# Uncovering the Mechanistic Landscape of Regulatory DNA with Deep Learning

**DOI:** 10.1101/2025.10.07.681052

**Authors:** Evan E. Seitz, David M. McCandlish, Justin B. Kinney, Peter K Koo

## Abstract

The regulatory genome encodes the logic that governs gene expression, enabling cells to respond to developmental, environmental, and evolutionary cues. This logic arises from complex *cis*-regulatory mechanisms that integrate transcription factor motifs, their syntactical arrangement, and surrounding sequence context, features that remain challenging to decode. Here, we present SEAM (Systematic Explanation of Attribution-based Mechanisms), a computational framework that combines deep learning with explainable AI to map the mechanistic impact of genetic mutations. Applied to human and *Drosophila* regulatory loci, SEAM uncovers functional binding sites at sequences of interest and identifies which mutations preserve, disrupt, or create novel binding sites. SEAM also reveals that two qualitatively distinct classes of regulatory signal are operative at many loci: signals that are robust to mutation and signals that are readily reprogrammable. These results clarify the inherent ability of regulatory DNA to evolve. They also position SEAM as a versatile framework for interpreting non-coding variants and for informing the mechanism-aware design of synthetic sequences.

---

Understanding how regulatory DNA controls gene expression remains one of the central challenges in biology. The regulatory genome exhibits a rich combinatorial architecture in which transcription factor binding sites (TFBSs)—short DNA motifs recognized by transcription factors (TFs)—are embedded within a broader sequence context shaped by nucleotide composition, simple sequence repeats, motif spacing and orientation, and local chromatin state.^1–7^ At any given regulatory locus, the precise combination and arrangement of these elements in a particular chromatin and cell state specify a *cis*-regulatory mechanism: a configuration of TFBSs and contextual interactions that determine transcriptional output. Mutations can disrupt or strengthen TF binding, create *de novo* sites, or alter motif syntax and TF cooperativity,^8–10^ reshaping regulatory logic in ways that drive evolutionary adaptation, disease susceptibility, or loss of normal function.^11–17^ Deciphering these mechanisms is essential for linking genotype to phenotype, revealing how regulatory programs shape development and evolution, and enabling the rational design of sequences for therapeutic and biotechnological applications.

Deep neural networks (DNNs) have become powerful tools for modeling *cis*-regulatory elements, enabling accurate prediction of regulatory activity, such as chromatin accessibility and gene expression, directly from DNA sequence.^18–22^ Despite their predictive power, DNNs remain difficult to interpret, earning them a reputation as a “black box.” *Post hoc* attribution methods—including *in silico* mutagenesis (ISM),^23^ Saliency Maps,^24^ and DeepSHAP^25^—quantify the contribution of each nucleotide to a model’s prediction, producing an attribution map that highlights the sequence features underlying the output (Fig. 1a,b).^26,27^ While these maps have proven the ability to reveal complex regulatory features, such as motifs, their syntax, and integration within sequence context,^28–33^ they reduce model interpretation to per-nucleotide importance scores. Consequently, they miss higher-order dependencies and provide little insight into the broader *cis*-regulatory mechanisms captured by the DNN, or how those mechanisms are rewired by mutations (Fig. 1c).

**Figure 1.**
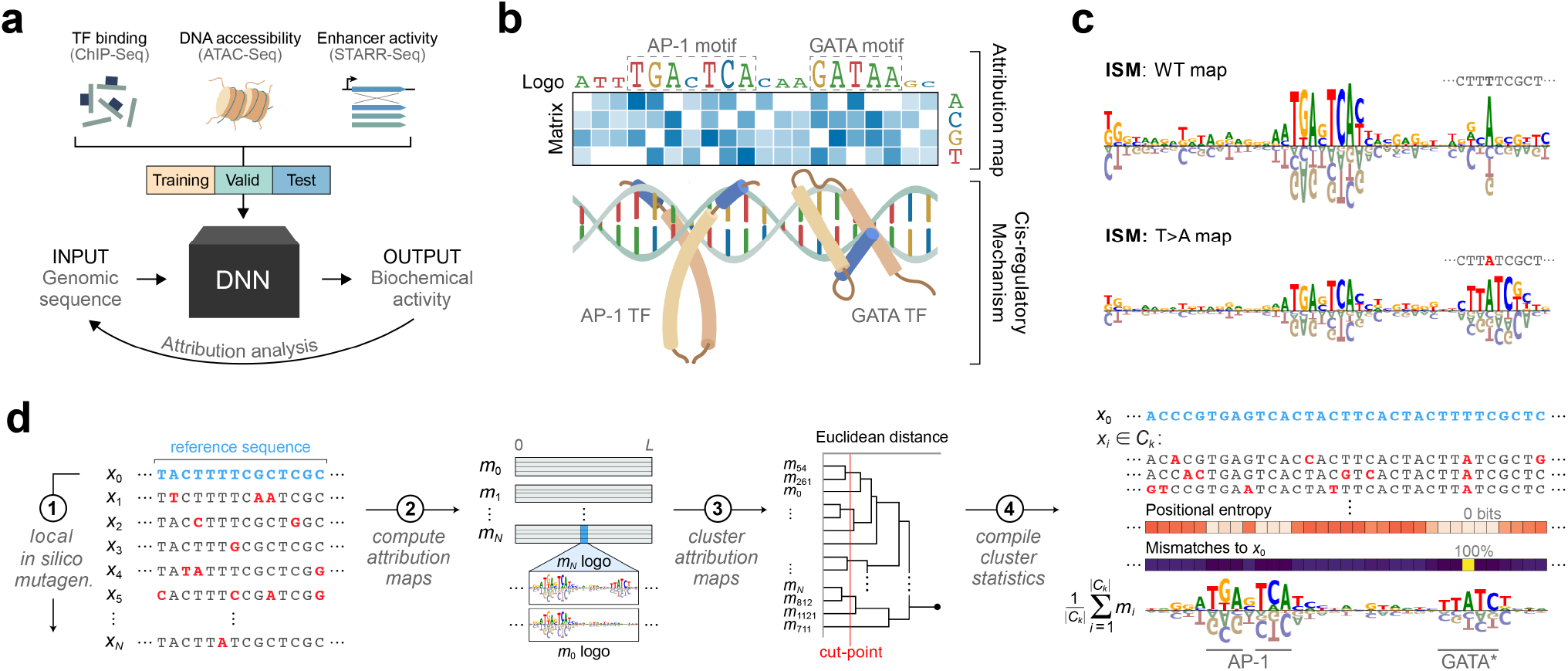
Overview of SEAM. **a**, Schematic of a deep learning workflow with attribution analysis. **b**, Schematic illustrating the relationship between attribution maps and *cis*-regulatory mechanisms which often represent protein-DNA interactions. **c**, Sequence logos for an *in silico* mutagenesis map of a regulatory sequence before (top) and after (bottom) a point mutation (shown in red). **d**, Schematic of the SEAM framework comprising four main steps: (1) generate an *in silico* sequence library; compute attribution maps for every sequence in the library using a genomic DNN; (3) cluster attribution maps to reveal distinct mechanisms; and (4) calculate statistics from each cluster to uncover sequence determinants driving mechanistic variation.

Here, we introduce SEAM (**S**ystematic **E**xplanation of **A**ttribution-based **M**echanisms), a computational framework that integrates deep learning, virtual perturbations, and explainable AI to reveal sequence-mechanism relationships: the sequence determinants that shape *cis*-regulatory mechanisms learned by a genomic DNN and how these mechanisms are rewired by specific mutations. Applied across diverse models, organisms, and functional genomics tasks, SEAM uncovered latent regulatory motifs, decomposed *cis*-regulatory mechanisms into the contributions of specific binding sites and their surrounding context, and revealed how small numbers of mutations can reprogram regulatory logic. These analyses highlight the remarkable evolvability of regulatory DNA, where minimal sequence changes can reconfigure motif syntax or activate new mechanisms. Together, SEAM establishes a general framework for moving beyond nucleotide-level importance toward mechanistic interpretation, enabling systematic dissection of regulatory logic, principled interpretation of noncoding variants, and rational, mechanism-aware design of synthetic regulatory elements.

## Results

### SEAM: Decoding the mechanistic impact of genetic variation

SEAM provides a computational framework for decoding the mechanistic impact of genetic variation within defined regions of sequence space (Fig. 1d). The workflow proceeds in four steps. First, SEAM generates a synthetic sequence library by partially mutagenizing a reference sequence, systematically sampling local sequence variants anchored to a common alignment. Second, it computes attribution maps for each sequence using a trained sequence-to-function DNN, translating model predictions into interpretable mechanisms. Third, these attribution maps are clustered to group sequences that share regulatory mechanisms, with averaging of attribution maps within clusters to reduce noise and emphasize consistent patterns. Finally, the clustered sequences are analyzed to identify regulatory elements, reveal how mutations rewire regulatory logic, and highlight potential routes of regulatory innovation.

A key design choice is the restriction to local sequence neighborhoods through partial random mutagenesis. By introducing only limited numbers of mutations, SEAM ensures that mechanisms remain anchored to a common reference, enabling whole-map clustering and preserving the full motif syntax. This contrasts with approaches such as TF-MoDISCo,^29^ which cluster short subsequences (“seqlets”) extracted from attribution maps for motif discovery, but cannot capture higher-order syntax across entire loci. Because SEAM analyzes aligned variants within a neighborhood, it allows systematic dissection of motif robustness, positional precision, and genetic basis. Moreover, by calculating attribution maps for mutated sequences, SEAM inherently incorporates higher-order interactions between *cis*-regulatory features learned by genomic DNNs, revealing the specific mutations that drive mechanistic changes.

SEAM is implemented as FAIR (Findable, Accessible, Interoperable, and Reusable) software and is designed to be both flexible and extensible. It accommodates a range of attribution methods, clustering strategies, and sequence library designs, and already encompasses several alternative strategies. Unless otherwise noted, we applied it using libraries of 100,000 variants generated by applying 10% random mutagenesis to a seed genomic sequence under investigation, DeepSHAP^25^ for attribution maps, and hierarchical clustering.^34^ Full implementation details and alternative options are provided in the Methods, whereas conceptual distinctions from related approaches are described in Supplementary Note 1.

### SEAM dissects complex *cis*-regulatory mechanisms

To demonstrate SEAM’s ability to resolve mechanisms within *cis*-regulatory sequences, we applied it to DeepSTARR,^28^ a sequence-to-activity DNN trained to predict enhancer activity measured using UMI-STARR-seq (Unique Molecular Identifier–Self-Transcribing Active Regulatory Region sequencing) in *Drosophila* S2 cells (see Methods). Previous analyses of DeepSTARR combined attribution maps with *in silico* perturbations to reveal dependencies between core TFBSs and their flanking nucleotides, as well as distance-dependent cooperative interactions between pairs of motifs.^35^ Several of these mechanisms were validated experimentally.^28,36^ While high-activity enhancers typically exhibit clear motif-based signals in attribution maps, sequences spanning the full activity range often show diffuse or obscured patterns that complicate the reliable identification of putative TFBSs. For example, some enhancers exhibit dense clusters of attribution scores, suggestive of overlapping or closely spaced motifs (Fig. 2a).

**Figure 2.**
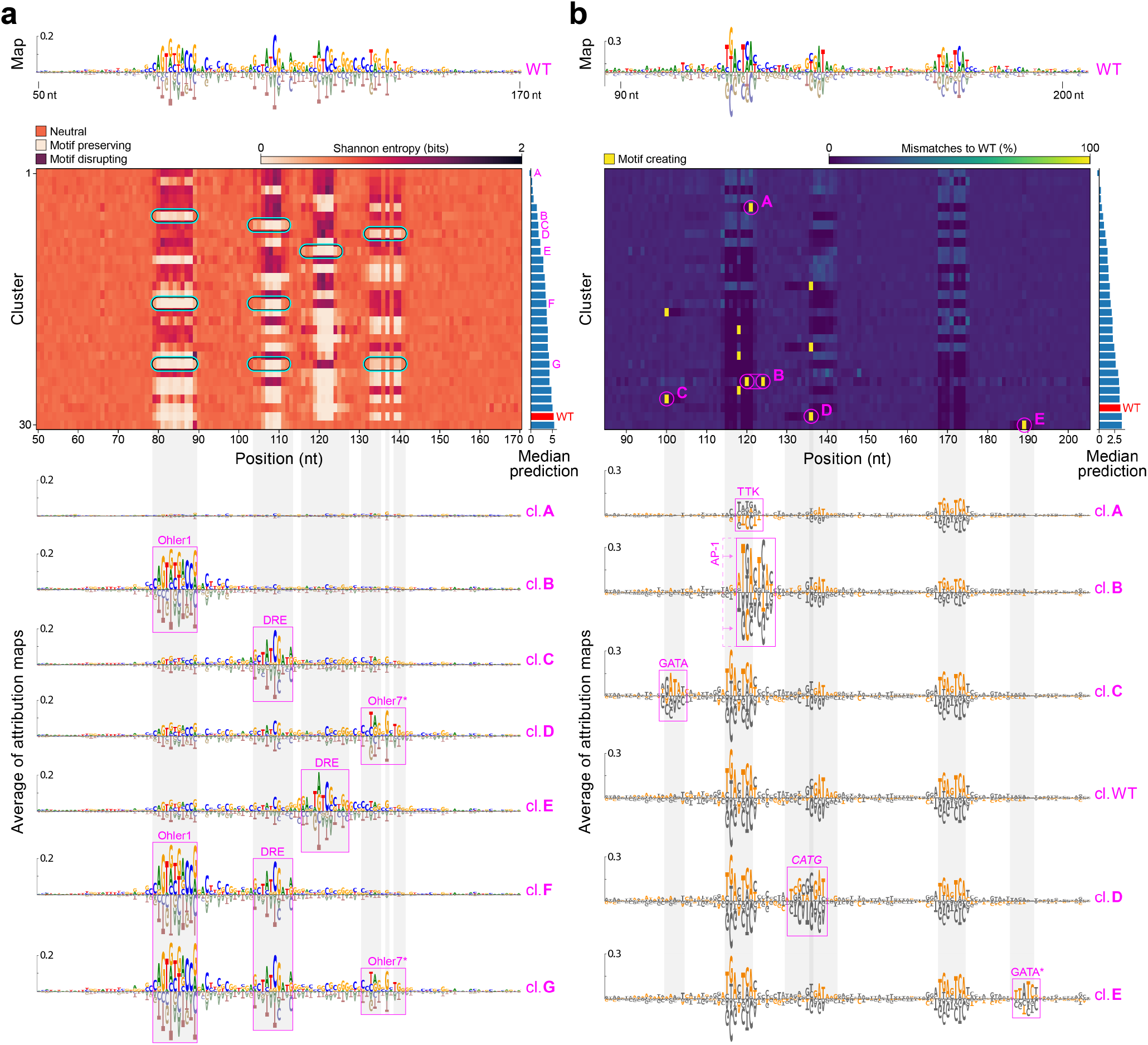
SEAM analysis of DeepSTARR mechanisms reveals active and poised regulatory motifs. **a**, Top: Attribution map for a wild type (WT) sequence obtained from the DeepSTARR test set (index 20647) using the housekeeping head. Middle: CSM-entropy generated using a partial mutagenesis library centered on the WT sequence, where clusters are ordered by their median DeepSTARR prediction. Each row represents a different cluster. The background entropy is ∼0.63 bits. Bottom: Sequence logos for various cluster-averaged attribution maps. **b**, Similar analysis performed on another enhancer locus (index 22612) using DeepSTARR’s developmental head with the exception that the CSM is based on percent mismatches between sequences within a cluster and the WT sequence. Bottom, sequence logos colored according to the WT sequence. **a**,**b**, Gray vertical bars represent high entropy regions shared across clusters in the CSM.

SEAM enables systematic dissection of these complex regulatory mechanisms. After clustering the attribution maps of mutagenized sequences, SEAM summarizes the distribution of mutations within each cluster using a Cluster Summary Matrix (CSM), where rows correspond to clusters, columns to nucleotide positions, and each entry is colored by a position-specific statistic (see Methods). For example, a CSM constructed based on the Shannon entropy of nucleotide frequencies (CSM-entropy) reveals striking patterns across clusters (Fig. 2a). In particular, entropy values can be grouped into three main categories: (1) *motif-preserving* positions which remain conserved across all sequences in a cluster (∼0 bits), indicating bases essential for that mechanism; (2) *motif-disrupting* positions, characterized by high entropy, where consistent disruption of a motif or feature defines the mechanism; and (3) *neutral* positions, showing background-level entropy (∼0.63 bits for a 10% mutation rate) that does not influence cluster assignment.

Vertical bands in the CSM-entropy highlight positions where motifs are present in the wild-type sequence, with entropy values indicating whether these motifs are preserved or disrupted. Contiguous motif-disrupting signatures suggest that multiple alternative mutations, or combinations thereof, can destroy a binding site. Occasionally, SEAM also detects motif-preserving signatures that arise outside these vertical bands, representing a subclass of motif-preserving behavior – the emergence of *de novo* motifs (Supplementary Fig. 1a). In this way, SEAM distinguishes positions that are mechanistically inert from those that drive regulatory change.

Averaging attribution maps within each cluster and visualizing them as sequence logos reveals a clear correspondence between CSM-entropy patterns and the presence or absence of motifs in that cluster’s attribution maps (Fig. 2a, bottom). Covariance analysis of the CSM-entropy allows precise segmentation of motifs and quantifies the co-occurrence of combinatorial motif states (Supplementary Fig. 2a-e). Further, motif epistatic interactions can be quantified by comparing predicted activities across clusters defined by distinct combinatorial states (Supplementary Figs. 2f,g; see Methods).

To gain additional insights, we constructed CSMs based on the percent mismatch of each cluster’s sequences relative to the wild type (CSM-mismatch, see Methods). Interestingly, a subset of clusters contained a specific point mutation(s) that consistently appeared in all sequences assigned to the cluster (Fig. 2b, Supplementary Fig. 3). These mutations corresponded to the creation of *de novo* binding sites, which were also visible in the entropy-based CSM (Supplementary Fig. 1a). For example, a single-nucleotide mutation converted an activating AP-1 motif into a TTK repressor site, producing a mechanism switch that drastically reduced predicted activity (Fig. 2b, cluster A). SEAM also identified a pairwise mutation that shifted an AP-1 motif three nucleotides to the right while maintaining activity similar to wild type (Fig. 2b, cluster B), suggesting an alternative binding configuration preserved by balancing selection. In addition to motif-creating mutations, the CSM-mismatch exposed multiple single-nucleotide substitutions in the core of an Ohler1 motif that fine-tuned activity across a dynamic range (Supplementary Fig. 3b). Thus, CSM-mismatch provides a powerful way to pinpoint specific mutations that reprogram *cis*-regulatory mechanisms.

**Figure 3.**
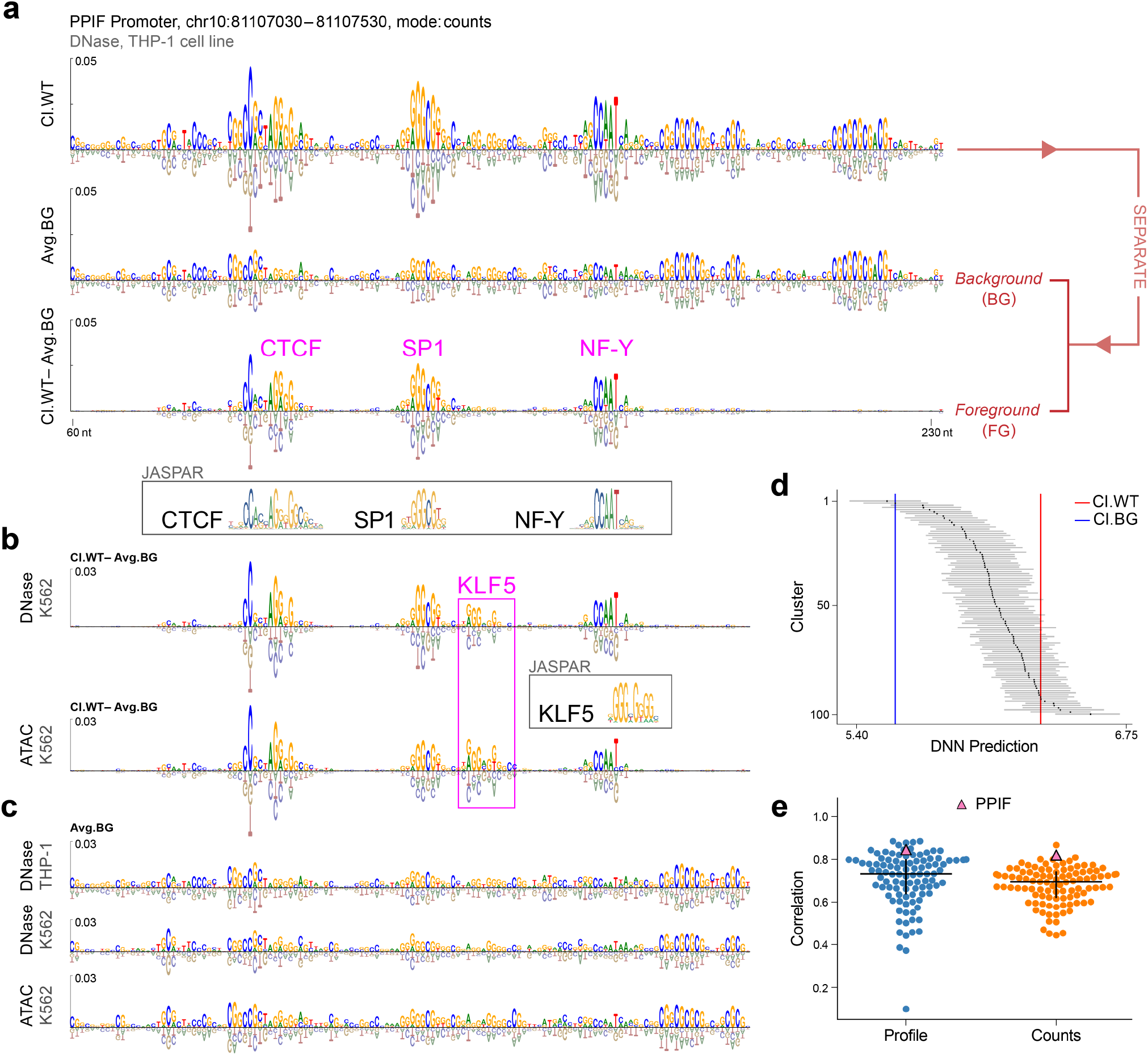
SEAM decomposes attribution signals according to mutational sensitivity. **a**, Top: Cluster-averaged attribution map for the WT cluster (cl. WT) at the PPIF promoter, obtained from a ChromBPNet model trained on DNase-seq readouts in THP-1 cell lines. Middle: Attribution map of the background attribution signal (BG), defined by the intra-cluster robustness to partial random mutagenesis (Avg. BG). Bottom: Attribution map of foreground attribution signal (FG), defined by the removal of BG from a given cluster (here, from the WT cluster). The resulting motifs in the foreground closely match the associated JASPAR entries: MA0139.1, MA2279.1.rc, MA0060.1. **b**, Foreground attribution comparison for the same PPIF region from ChromBPNet models trained independently on K562 using DNase-seq (top) and ATAC-seq (bottom). Foreground attributions in K562 contain an additional KLF5 motif (JASPAR entry MA0599.1.rc) compared to THP-1. **c**, Comparison of the background attribution maps for the same PPIF region for the three ChromBPNet models independently trained on different assays and cell types. **d**, Box plots showing DNN predictions for each SEAM-derived cluster for ChromBPNet trained on DNase with THP-1. Lines representing the upper and lower quartiles. **e**, Swarm plot of the Pearson correlation between SEAM background attribution maps from paired ChromBPNet models trained on different data modalities (DNase-seq vs ATAC-seq in K562) for the two model outputs (profile head and counts head). Each point represents a different promoter.

These findings were robust across different genomic DNNs trained on different experimental assays and biological systems (Supplementary Fig. 1), as well as SEAM parameter choices, including the number of clusters (Supplementary Fig. 4), clustering algorithms (Supplementary Fig. 5), mutation rates and library sizes (Supplementary Fig. 6), and choice of attribution methods (Supplementary Fig. 7). Together, these results demonstrate SEAM’s ability to reliably resolve the diversity of *cis*-regulatory syntax and its impact on regulatory activity.

**Figure 4.**
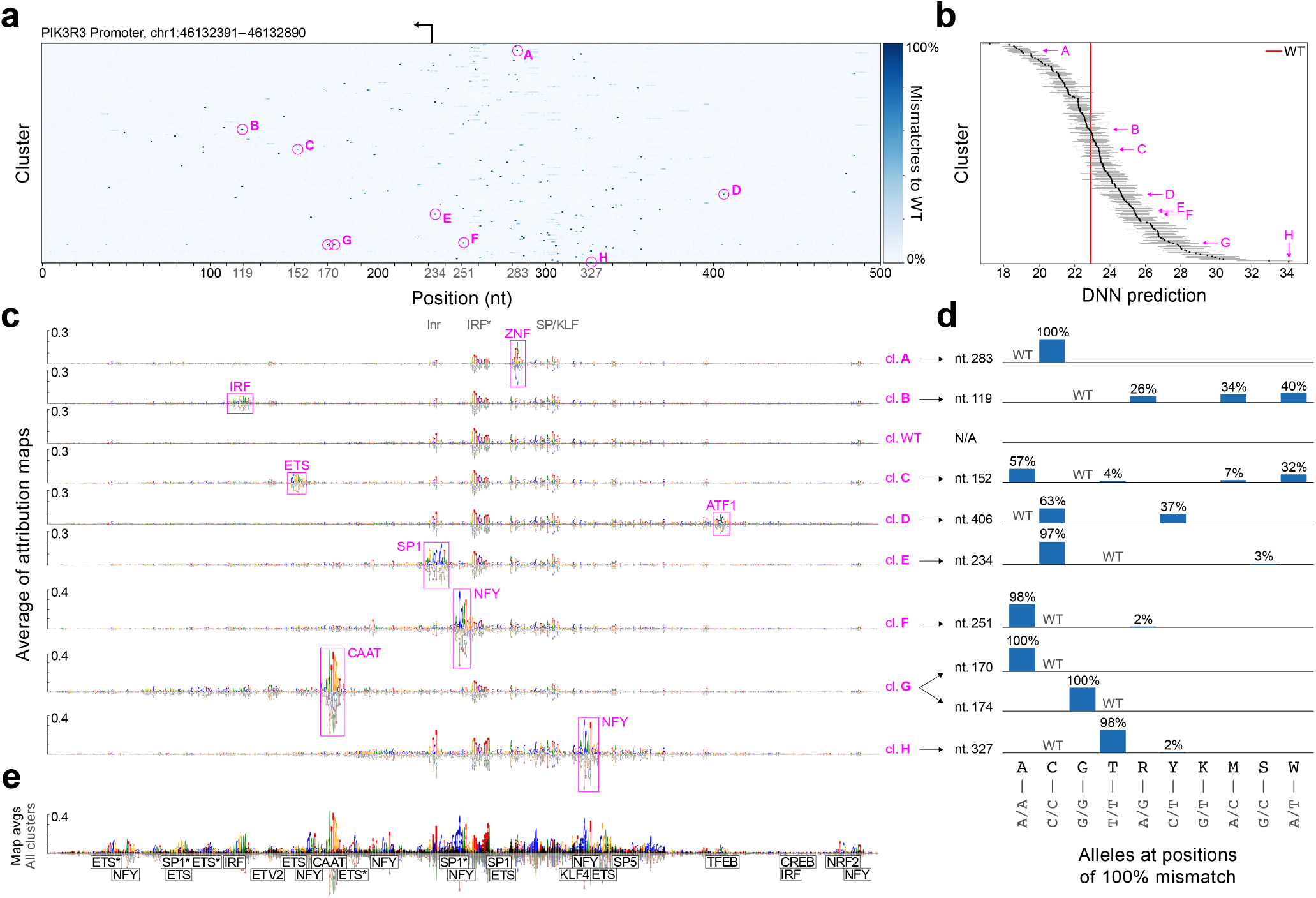
SEAM analysis of CLIPNET PIK3R3 promoter reveals the high evolvability of regulatory motifs. **a**, CSM-mismatch for the PIK3R3 promoter, centered at the TSS, colored by percent mismatches to the WT sequence for each of the 200 clusters. **b**, Box plots showing DNN predictions for each cluster (matched rows as CSM), with lines representing the upper and lower quartiles. **c**, Sequence logos of foreground attribution maps for select clusters. Pink boxes annotate the presence of a *de novo* regulatory motif. **d**, Bar plot of nucleotide substitution frequencies, based on IUPAC codes, at the position of the 100% mismatch for sequences associated with each of the example clusters. **e**, Overlay of all 200 cluster-averaged attribution maps.

**Figure 5.**
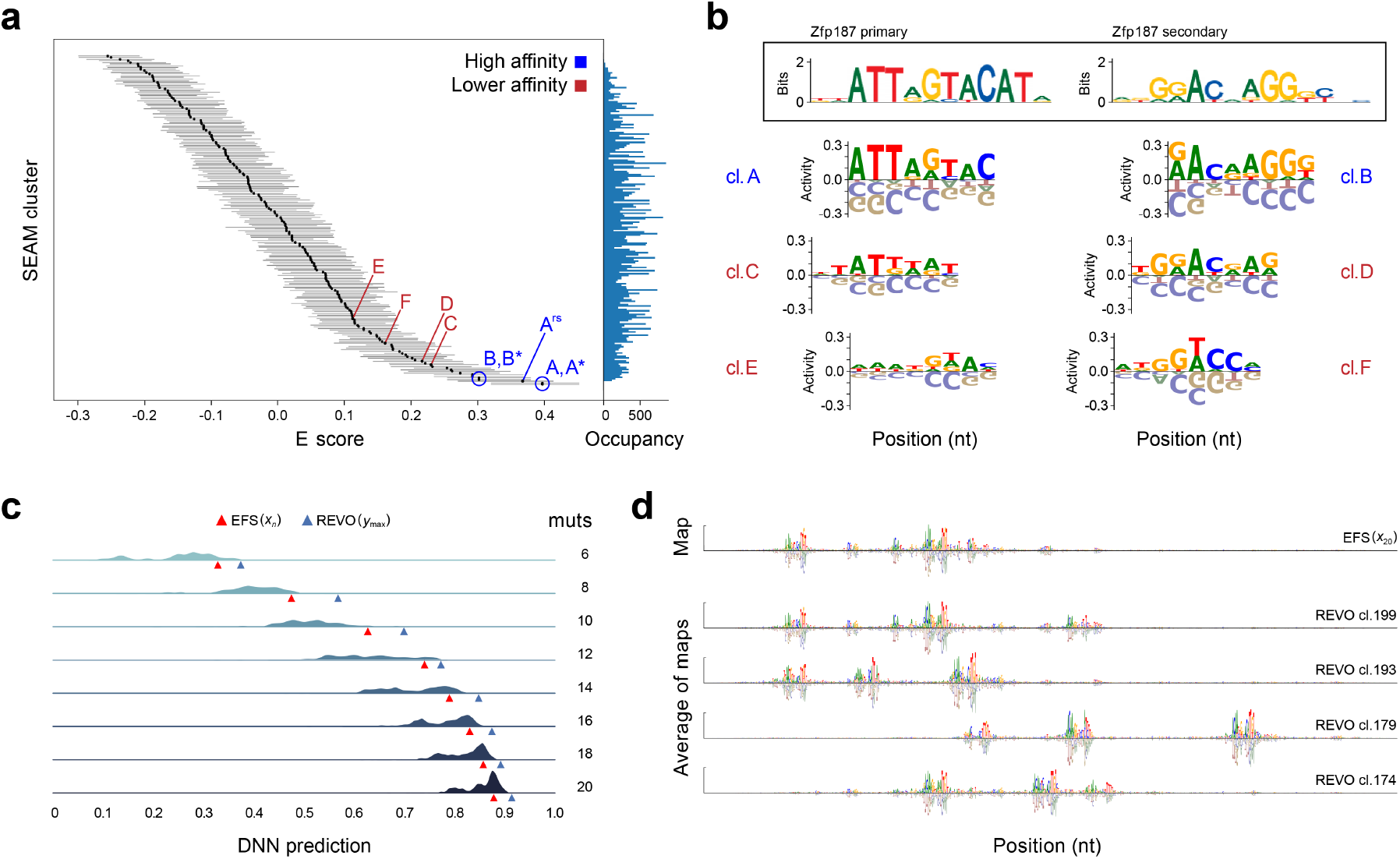
SEAM generalizes to diverse sequence libraries. **a**, SEAM analysis on the Zfp187 protein binding microarray data. Box plots show E scores for each of the 200 clusters. Lines represent the upper and lower quartiles; the average IQR across clusters is 0.12. Superscripts denote either a reverse complement (*) or a register shift (rs) of the same motif for that label. The plot on the right shows the occupancy (i.e., number of empirical mutagenesis maps) in each cluster. **b**, Top: Sequence logos for two alternative binding modes captured in the original PBM study using the Seed-and-Wobble algorithm. Bottom: Sequence logos of averaged empirical mutagenesis maps corresponding to SEAM clusters. **c**, Distribution of DeepMEL2 predictions for oracle-based designed sequences using a starting sequence EFS-6 (from the original analysis). The highest-scoring sequence in each distribution (blue triangle) surpasses the score of ISE (red triangle) at each mutational step. **d**, Top row: Attribution map for the ISM-designed sequence. Remaining rows: Mechanistically diverse attribution maps for REVO-designed sequences with high predicted activities.

### SEAM disentangles motif syntax from sequence context in regulatory mechanisms

In the course of SEAM analyses, we observed a surprising distinction between two types of attribution signals: cluster-specific patterns that appear to reflect TF binding sites, and diffuse locus-specific signals that remained largely constant across clusters. Whereas the TF binding site signals were readily disrupted by key mutations, the diffuse signals showed little or no change across sequence variants, indicating they reflect properties common to all sequences within that local region of sequence space. We term the variable motif syntax that reflects TF binding sites the *foreground* and the mutation-insensitive signal the *background*.

To isolate the background, we developed an information-based analysis (see Methods). We first applied this decomposition to ChromBPNet, a sequence-to-activity DNN trained to predict chromatin accessibility profiles and counts from DNase-seq data in the THP-1 human cell line.^20^ At the PPIF promoter, where ChromBPNet predictions have been shown to align with perturbation measurements from VariantEFFECTS,^10^ subtracting the background sharpened TF motifs and clarified motif syntax, yielding foreground maps with binding sites that closely matched known motifs (Fig. 3a, bottom; Supplementary Fig. 8). In this case, the background signal correlates with GC content (Fig. 3a, middle), consistent with the established role of GC content in mammalian nucleosome positioning and gene expression.^37–39^

We next asked whether backgrounds vary across loci and species. Indeed, both their nucleotide composition and relative strength differed markedly. At the PPIF enhancer, the background exhibited a more balanced profile across all four bases (Supplementary Fig. 9), whereas in *Drosophila* enhancers, backgrounds were often A/T-rich (Supplementary Fig. 10), consistent with the known role of poly-A/T tracts as TF recruitment scaffolds.^40^ The strength of the background also varied: in highly active sequences, foreground motifs stood out clearly, while in others, strong background signals obscured low-affinity or cryptic motifs. In such cases, background subtraction revealed these features with striking clarity (Supplementary Figs. 11 and 12).

To assess whether these backgrounds reflected technical or model-specific artifacts, we confirmed that they were consistent across ChromBPNet models trained on different data subsets and robust to changes in mutation rate (Supplementary Fig. 13). Backgrounds were also nearly identical across different attribution methods (Supplementary Fig. 14), indicating that they are not specific to a particular attribution technique.

To evaluate whether backgrounds reflect broader biological properties, we compared ChromBPNet models trained independently on ATAC-seq and DNase-seq data, two distinct assays of chromatin accessibility. Background attributions were highly concordant across modalities (Fig. 3c) and remained consistent across clusters even when motif syntax differed and accessibility levels varied (Fig. 3d). This correspondence was further quantified across additional genomic loci (Fig. 3e). As an illustrative example, the PPIF promoter, which is broadly expressed across cell types, showed strong correlations between Jurkat and K562 (Fig. 3c). Although the precise origins of these background signals remain unclear, their reproducibility across models, assays, and species suggests that they reflect underlying biological sequence properties rather than artifacts of the model, SEAM parameter choices, or technical biases in the assays.

Together, these results demonstrate that sequence context encodes an underappreciated layer of regulatory information, captured by genomic DNNs but only made accessible through SEAM’s foreground–background decomposition. Whereas standard single-sequence attribution produces overlapping signals that blur motif syntax and background context, SEAM disentangles these components by systematically probing local sequence space. In doing so, SEAM provides a framework for dissecting complex *cis*-regulatory mechanisms and offers a new lens for interpreting regulatory DNA.

### Regulatory sequences exhibit high mechanistic evolvability

SEAM exposes the remarkable evolvability of regulatory DNA, reveals that even single variants can reprogram regulatory mechanisms, including variants associated with disease. A substantial proportion of disease-linked variants lie in noncoding regions,^16^ including introns and promoters, where they can act by altering or creating TFBSs.^17^ To investigate how specific alleles reprogram transcriptional regulation at polymorphic loci, we applied SEAM to CLIPNET, a sequence-to-activity DNN trained on heterozygous encodings to predict PRO-cap (Precision Run-On and sequencing of capped RNA) data across individuals in human lymphoblastoid cell lines.^41^ As a case study, we focused on the promoter of *PIK3R3*, a highly polymorphic locus involved in diverse cellular processes and notably overexpressed in cancer.^42–44^

Using a 1% partial random mutagenesis library, SEAM uncovered a striking diversity of transcriptional activities and associated mechanisms (Fig. 4), with many novel regulatory programs driven predominantly by single nucleotide mutations (Fig. 4a, cluster A) that quantitatively tune transcriptional activity (Fig. 4b). For example, SEAM identified an activity-reducing mutation that generates a *de novo* zinc finger motif near the transcription start site (Fig. 4c; cl. A). Conversely, motifs required for transcription initiation, such as the initiator element (Inr), were largely preserved across most clusters. However, in some cases, conserved elements could be entirely overwritten by a few mutations, yielding alternative regulatory logic. In one instance, a pairwise mutation created a CAAT box that upregulated transcriptional activity and reversed the predicted direction of transcription (Fig. 4c, cluster G; Supplementary Figs. 15a,b). To further investigate this effect, we independently mutated each of the two nucleotides and examined the resulting attribution maps and model predictions for both the positive and negative ProCap strands (Supplementary Fig. 15a). Single mutations had little effect: the attribution maps were largely unchanged, and predicted transcriptional activities differed only modestly from the wild-type sequence. By contrast, mutating both nucleotides simultaneously generated a clear CAAT box motif and caused the initiator motif to shift from the right to the left of the newly formed CAAT box (Supplementary Fig. 15b). This led to a pronounced change in the predicted transcriptional activity and direction of transcription. Notably, the initiator motif was already present in the wild-type sequence but was predicted to be non-functional until a CAAT box was brought into close proximity. This finding demonstrates that CLIPNET is not merely detecting motifs in isolation, but is also sensitive to the motif syntax when assigning functional importance.

While some mechanisms were defined by a single critical allele, others arose from sets of interchangeable alleles or strictly heterozygous configurations (Fig. 4d). This allele-specific mechanistic heterogeneity is difficult to resolve using conventional interpretability methods and underscores the high evolvability of the PIK3R3 promoter, where minimal sequence changes can activate distinct *cis*-regulatory programs.

Similar reprogramming potential was observed across additional human promoters, underscoring their intrinsic evolvability (Supplementary Fig. 16). Strikingly, SEAM revealed that oncogene promoters such as *MYC* and *NOTCH1* displayed greater mechanistic evolvability than other promoters, exceeding even those with comparable levels of polymorphism (Supplementary Fig. 16).

To further investigate the mechanistic complexity of promoters, we turned to ProCapNet, another DNN trained to predict PRO-cap data.^45^ Whereas CLIPNET enabled the study of allele-specific rewiring across individuals, ProCapNet was originally designed to dissect the regulatory logic of transcription initiation. At the *MYC* promoter, ProCapNet revealed that initiation is controlled by cooperative and competitive dependencies between motifs and TSSs, with strong interactions between TATA and Inr motifs, auxiliary contributions from SP1/BRE motifs, and asymmetric competition between adjacent sense and antisense TSSs.^45^

Applying SEAM to the *MYC* promoter, we recovered these previously described dependencies, but also revealed a broader mechanistic landscape by probing local sequence space. SEAM uncovered cryptic sense and antisense TSSs, alternative motifs such as TATA boxes and BRE/SP sites, and mutations that relocated TSSs or reversed transcriptional direction (Supplementary Fig. 17). Thus, SEAM complements motif ablation–based analyses by systematically charting the repertoire of initiation mechanisms encoded in promoter sequence space and accessible through just a few specific mutations.

### SEAM generalizes across diverse sequence libraries

Sequence space is vast, and no single library design can capture all of its mechanistic possibilities. SEAM is not universally applicable, but when applied to assays that generate libraries amenable to clustering, it can reveal distinct facets of regulatory logic. To demonstrate this, we analyzed three complementary regions of sequence space: (i) exhaustive mutagenesis libraries that expose the full spectrum of binding preferences, (ii) optimized libraries from directed evolution–based approaches that highlight alternative high-functioning solutions, and (iii) global libraries that probe broad properties of motifs and their context dependence. Together, these applications illustrate how SEAM extracts diverse mechanistic insights depending on the sequence space under study.

First, we applied SEAM to experimental combinatorial-complete mutagenesis libraries that probe all possible short sequences, providing a comprehensive view of binding preferences. Protein Binding Microarrays (PBMs), which measure TF enrichment scores across all *k*-mers, exemplify this type of dataset (Supplementary Fig. 18a).^46^ Applied to PBM 8-mer data for ZFP187, SEAM recovered both primary and secondary binding motifs (Fig. 5a,b). These results matched known preferences and further uncovered lower-affinity mechanisms often overlooked by conventional methods such as Seed-and-Wobble.^46^ Extending the analysis to Hnf4a, SEAM revealed a continuous landscape of binding strengths and syntactic variation by embedding attribution maps prior to clustering (Supplementary Fig. 18b–e). Thus, SEAM systematically organizes exhaustive libraries to resolve both strong and subtle binding preferences, exposing mechanistic diversity in TF recognition.

Next, we applied SEAM to sequence design libraries generated by iterative optimization algorithms such as *in silico* evolution (ISE). These approaches search sequence space for sequences with high regulatory activity, typically using local search strategies that converge on greedy solutions and fail to capture the diversity of possible regulatory mechanisms.^47–50^ To broaden this design space, we developed Redirected Evolution (REVO), a motif-centric extension of ISE that systematically blocks mutagenesis iteratively in specific high-attribution motifs before re-optimizing, thereby steering the search toward alternative regulatory solutions (see Methods; Supplementary Fig. 19a). Applying SEAM to REVO-derived libraries generated with DeepMEL2, a DNN trained to predict chromatin accessibility in melanoma,^49,51^ revealed a much broader repertoire of mechanisms than standard ISE. SEAM identified clusters defined by diverse motif types and configurations, many of which achieved activities comparable to or exceeding those of ISE-optimized sequences (Fig. 5c,d, Supplementary Fig. 19b). Together, these results show that REVO diversifies sequence optimization by uncovering alternative motif grammars, while SEAM organizes this diversity into interpretable mechanistic classes, enabling both broader exploration and clearer interpretation for rational regulatory sequence design.

We also extended SEAM to ‘global’ libraries, which anchor motifs within diverse sequence contexts and thereby provide a way to probe how context influences regulatory activity (Supplementary Note 2). Although these analyses are exploratory, they illustrate how SEAM can be applied beyond local mutagenesis and design-derived searches, highlighting its potential to dissect mechanisms across different regions of sequence space. Together, these applications demonstrate SEAM’s versatility in revealing the range of *cis*-regulatory logic accessible through complementary experimental and computational designs.

## Discussion

SEAM enables systematic dissection of *cis*-regulatory mechanisms by probing defined regions of sequence space through the lens of deep learning models. Rather than focusing on single sequences or isolated motif occurrences, SEAM organizes populations of related variants into clusters representing shared molecular mechanisms and pinpoints the mutations that drive transitions between these clusters. A central innovation is SEAM’s ability to disentangle complex attribution signals into two complementary components: a reprogrammable foreground signal, reflecting TF binding sites and motif syntax, and a mutation-insensitive background signal, reflecting stable sequence properties shared within a localized region of sequence space. Applied across diverse models and datasets, SEAM consistently shows that regulatory logic can be rewired through a few key mutations, underscoring the remarkable flexibility of regulatory DNA.

Because SEAM maps the alternative regulatory mechanisms accessible within a few mutations of the wild-type sequence, our analysis has particular relevance from an evolutionary perspective. For instance, SEAM readily identifies regulatory sequences that the DNN predicts are a single mutation away from forming a new binding site (e.g. Fig. 2b); such sequences have been previously discussed in the population genetics literature as critical for the evolution of novel binding sites on evolutionary time scales short enough to explain differences between sister taxa.^52^ Similarly, in many abstract models of evolution under complex genotype-phenotype maps, populations diffuse over large networks of fitness-neutral genotypes sharing the same phenotype until they find key ‘portal’ mutations that generate a novel beneficial phenotype.^11^ SEAM’s clusters can be viewed as providing a systematic catalog of these networks in sequence space, while also identifying the specific mutations that convert one mechanism into another (Supplementary Fig. 20).

A key strength of SEAM is its ability to disentangle foreground and background attribution signals. Foreground signals capture the effects of discrete TF binding sites, whereas background signals reflect diffuse, context-dependent patterns that remain stable in the face of mutations. Often dismissed as noise, SEAM shows that these background signals can be consistent across assays and thus likely reflect biologically meaningful signals. Indeed, the presence of these background signals may help explain why identical motif syntax can yield different outcomes in distinct sequence contexts. The mechanistic bases for these background signals remain uncertain, but they may reflect intrinsic DNA properties such as shape or flexibility, or epigenetic features that sequence-based models must implicitly infer.^7,53–56^ Clarifying these origins will be important for understanding how contextual features influence TF compatibility and why certain motif configurations are effective only in specific sequence environments.

Although our analyses here focus on regulatory DNA, SEAM is inherently model- and modality-agnostic. In this study, we used attribution maps from genomic DNNs as a proxy for regulatory mechanisms, reflecting the logic those models learn to connect sequence with activity. The same framework, however, can be extended to any sequence-function mapping, spanning RNAs, protein sequences and even cis-regulatory elements,^57^ provided an appropriate mutagenesis strategy and predictive model. Moreover, SEAM can operate directly on dense mutagenesis datasets, enabling model-free analysis of regulatory mechanisms when dense sequence–function measurements are available. This flexibility highlights SEAM’s broader potential to bridge AI-based modeling with experimental data across diverse biological domains.

SEAM is best understood as a meta-explanation framework—an explanation of explanations. Whereas methods such as TF-MoDISco cluster short segments of attribution maps (“seqlets”) to discover motifs from observed genomic sequences,^29^ SEAM instead clusters entire attribution maps, primarily from synthetic sequence libraries, to capture how binding sites and their contexts reconfigure across local sequence neighborhoods. This broader scope allows SEAM to dissect motif syntax and context dependencies, though it introduces limitations. Scalability depends on choices of library design, attribution method, and clustering strategy, which may need to be tuned to the application. In addition, because SEAM builds on attribution maps, it inherits their additivity assumption and can overlook higher-order dependencies. Nonetheless, SEAM naturally extends to richer attribution methods, including second-order analyses such as pairwise *in silico* mutagenesis^58,59^ and saliency-based interactions (e.g., Jacobians and Hessians),^60–62^ providing a path to capture more complex regulatory logic.

SEAM is released as an open-source, FAIR software framework for leveraging genomics AI to study *cis*-regulatory mechanisms. It clarifies motif syntax, provides tools to dissect motif–context dependencies, and distinguishes neutral from functional variation. More broadly, SEAM advances regulatory genomics toward mechanism-level reasoning. By systematically analyzing how mutations rewire regulatory logic, SEAM provides a conceptual and computational foundation for linking AI-based predictions to experimentally testable hypotheses and for guiding the rational design of regulatory DNA.

## Methods

### SEAM framework

SEAM is a computational framework for mapping the repertoire of *cis*-regulatory mechanisms within a population of sequences and identifying the mutations that drive mechanistic diversity. At a high level, SEAM first infers the regulatory mechanism underlying each sequence’s activity through attribution analysis, then groups sequences into clusters based on shared mechanistic features. Comparing clusters reveals how specific mutations reprogram regulatory logic and uncovers the sequence determinants of mechanistic shifts.

*Cis*-regulatory sequences are highly diverse, and the mechanisms that govern their function can differ substantially, even among elements with similar activity.^10^ This diversity makes it difficult to identify generalizable regulatory principles, particularly because sequences drawn from across the genome cannot be meaningfully aligned or clustered: they may encode entirely different architectures with no shared anchors. SEAM addresses this by restricting analysis to localized sequence neighborhoods, variant libraries generated by introducing limited mutations around a common reference sequence. Within such neighborhoods, sequences remain aligned, enabling SEAM to cluster attribution maps and isolate mechanistic differences that can be directly attributed to specific nucleotide substitutions. This local focus ensures that the analysis reveals genuine rewiring of regulatory logic, rather than artifacts of unrelated sequence architectures.

#### SEAM overview

The SEAM framework consists of four main steps.

1. **Variant library construction.** Starting from a reference sequence, SEAM generates a local sequence neighborhood by introducing partial random mutations at a specified mutation rate. This ensures that all sequences remain aligned and that mechanistic changes can be attributed to specific substitutions. Libraries can also be supplied directly from experimental or computational designs.
2. **Attribution map generation.** Each sequence in the library is scored with a trained sequence-to-function DNN, and attribution maps are computed to quantify the contribution of each nucleotide to predicted activity.
3. **Clustering attribution maps.** Attribution maps are compared and clustered, grouping sequences that are predicted to share regulatory mechanisms. Averaging maps within each cluster reduces noise and reveals the core mechanistic features underlying that group.
4. **Mechanistic interpretation.** SEAM computes sequence summary statistics across clusters—such as entropy- or mismatch-based Cluster Summary Matrices—to highlight motif-preserving, motif-disrupting, or motif-creating positions.

#### Cluster Summary Matrices

To summarize mutational patterns within SEAM-defined clusters, we constructed Cluster Summary Matrices. A CSM is a two-dimensional representation where rows correspond to clusters and columns correspond to nucleotide positions in the reference sequence. Each entry encodes a position-specific statistic computed over all sequences assigned to that cluster.

- **CSM-entropy**. For each cluster and nucleotide position, we calculated the Shannon entropy of the nucleotide distribution:

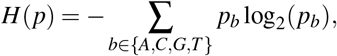

where *p*_*b*_ is the frequency of base *b* at that position among sequences in the cluster. Entropy values close to 0 indicate conservation (motif-preserving positions), while high entropy values indicate variability (motif-disrupting positions). Positions with entropy near the background expectation (approximately 0.63 bits for a 10% mutation rate) were considered neutral. The resulting CSM-entropy provides a positional map of conserved, disrupted, and neutral sites across clusters.
- **CSM-mismatch**. As a complementary representation, we computed the percent mismatch relative to the reference sequence at each nucleotide position. For a given cluster and position, this statistic is defined as the fraction of sequences in the cluster that differ from the reference base. Unlike entropy, which reflects overall variability, the mismatch statistic highlights specific point mutations that consistently recur across a cluster. The resulting mismatch-based CSM enables identification of single-nucleotide substitutions or small sets of mutations that reprogram regulatory mechanisms.

#### SEAM outputs

SEAM produces five key outputs: (1) a set of distinct regulatory mechanisms sampled by the sequence library, including the assignment of each sequence to its corresponding cluster; (2) the predicted activity distribution for each cluster, summarizing the functional variation among sequences with similar mechanisms; (3) an averaged attribution map for each cluster, capturing the shared regulatory features across sequences with similar mechanisms; (4) the CSM, which reports position-wise nucleotide frequencies across sequences in each cluster, enabling entropy-based or mismatch-based assessments of sequence variability; and (5) background-separated attribution maps, which disentangle mutationally-sensitive and -insensitive attribution signals and reveal a distinct background signal that is itself biologically interpretable. Together, these outputs provide a versatile set of resources for investigating the functional and mechanistic variation within regulatory sequences.

While this description is tailored for applying SEAM on a “local” sequence library, SEAM is a flexible framework that can be customized through the selection of different mutagenesis strategies (see “SEAM libraries”, below) for various regulatory genomics applications.

### Attribution methods

Our analyses used attribution maps computed using a variety of methods, implemented as follows.

- ***In silico* Mutagenesis** was computed by evaluating the change in the scalar DNN prediction from the WT prediction for every single nucleotide variant in a given sequence.^23^
- **Empirical Mutagenesis** used experimental measurements in place of DNN predictions. In the case of PBM data, for every sequence in the combinatorial-complete library, SEAM generates an empirical mutagenesis map using the log2 fold change between the E scores of the reference sequence and each SNV.
- **Saliency Maps** were computed by evaluating the gradient of the scalar DNN prediction with respect to the one-hot encoding for a given sequence.^24^
- **SmoothGrad** maps were computed by averaging Saliency Maps for 50 noise-injected sequences.^63^ Each noisy sequence was computed by adding Gaussian noise (mean zero, standard deviation 0.25) to each nucleotide variant in the one-hot encoded sequence.
- **Integrated Gradients** (IntGrad) was computed by averaging attribution maps for sequences that are interpolated between a baseline reference sequence and the sequence of interest. The gradient of the DNN’s scalar prediction is integrated along the interpolated path, and the resulting attributions reflect the cumulative contribution of each nucleotide to the prediction.^64^ The zero baseline was chosen for all examples in this analysis.
- **DeepSHAP** scores were computed using the SHAP package^25^ or DeepLIFT package,^65^ with the same hyperparamters and package as the original study for each respective genomic DNN. We employed 100 dinucleotide-shuffled sequences was used as the baseline for all examples in this analysis.

All attribution methods were zero-centered along the channel axis as a correction,^66^ a ‘gauge fixing’ trick that ensures attribution maps better reflect interpretable *cis*-regulatory mechanisms.^67^

### Clustering methods

SEAM supports clustering of attribution maps either directly in the original high-dimensional space or after dimensionality reduction via embedding. Aligned attribution maps can be cropped around a region of interest prior to clustering, effectively modulating the resolution of mechanistic details uncovered.

- **Hierarchical clustering** was applied using Ward’s linkage on the Euclidean distance matrix computed from the attribution map library.^34^ This method iteratively merges clusters to minimize within-cluster variance, producing a dendrogram that groups attribution maps by similarity. The dendrogram is then cut at a specified level to define the final set of clusters. For hierarchical clustering, distance calculations were refactored within the SEAM API as a class-based object optimized for GPU-accelerated with memory-mapped batch processing to handle large datasets efficiently. Actual linkage calculations and cluster formations were implemented using the Scipy^68^ cluster.hierarchy package.
- **Other clustering**, such as K-means^69^ or DBSCAN^70^ clustering, can be performed directly on attribution maps or upon dimensionality reduction using principal component analysis.^71^ Visualization of K-means clusters can be through low-dimensional (non-linear) embeddings based on t-SNE^72^ or UMAP.^73^ Sklearn^74^ implementations were used for PCA, K-Means, and DBSCAN.

### SEAM libraries

SEAM is a versatile framework that can be applied to a range of sequence library designs, including but not limited to local, global, complete, and sequence design libraries.

- **Local library**. Random partial mutagenesis is applied to a genomic sequence of interest to sample *N* aligned synthetic sequences. The number of mutations in each individual sequence is a Poisson distributed random variable having mean *Lr*, where *L* is the sequence length and *r* is the mutation rate.
- **Combinatorial complete library**. A set of sequences that includes all possible combinations of nucleotides at specified positions, ensuring comprehensive coverage of sequence variation. In this analysis, SEAM was applied directly to experimental protein binding microarray datasets that met this requirement by converting the complete sequence-function dataset into a complete set of empirical mutagenesis maps.
- **Sequence design library**. In this study, we used REVO (see below) to generate a library of heterozygous sequences anchored at a genomic sequence of interest.
- **Global library**. Fixed genetic elements, such as putative TFBSs, are embedded at the same position across a library of random sequences.

### Sequence design methods

We generated sequence design libraries using two iterative optimization strategies: *in silico* evolution and Redirected Evolution. Both approaches start from a random, GC-adjusted sequence and evolve variants predicted to increase regulatory activity.

- ***In Silico* Evolution**. ISE is an established greedy *in silico* directed evolution algorithm.^47^ At each iteration, all possible single-nucleotide variants of the current sequence are scored with the model, and the SNV with the highest predicted increase in activity is selected. This process is repeated until the sequence accumulates *t* mutations, typically producing a single optimized sequence after 10–15 iterations. In this study, ISE was applied to DeepMel2 following their naming convention: Evolved from Scratch (EFS).^49^
- **Redirected Evolution**. REVO extends standard ISE by enforcing diversity across optimization trajectories. The procedure begins with a standard ISE run from a random, GC-adjusted seed sequence. After this run converges on an optimized sequence with *t* accumulated mutations, we compute an attribution map for the final sequence. Sliding windows of length *w*_*l*_ are moved across the attribution map, and each window is scored by summing the maximum attribution values at each position. The top *w*_*n*_ non-overlapping windows are retained as high-attribution regions that represent the dominant features of the optimized sequence. From these regions, REVO constructs *protection masks*. Each mask corresponds to one or more high-attribution regions that are held fixed (i.e., forbidden from mutation) in subsequent ISE runs.

For example, if three regions are selected, REVO generates new ISE runs where region A is protected, region B is protected, region C is protected, A+B are protected, and so on. Each mask defines a separate branch in a search tree. For each branch, ISE is restarted from the original random seed sequence, run again for *t* iterations, but now under the specified protection constraints – hence *redirected evolution*.

After each new branch completes its optimization, REVO recomputes the attribution map for that branch’s optimized sequence, identifies its dominant high-attribution regions, and generates new protection masks. This process is repeated for *T* rounds, producing a branching tree of optimization paths. Redundant branches that converge to the same optimized sequence are pruned in real time to avoid unnecessary computation. In this way, REVO iteratively redirects evolution away from previously discovered high-attribution motifs, forcing exploration of alternative motifs and configurations. All sequences generated across all branches are collected to form the REVO library, which is subsequently analyzed using SEAM. For the analyses in this study, we combined libraries across *w*_*l*_ ∈ {5, 10, 15, 20}, with *w*_*n*_ = 4, *t* = 20, and *T* = 3. All resulting sequences were used as inputs to SEAM.

### SEAM-based background separation

When using a local library, an approximately uniform set of backgrounds emerges in the attribution maps across all sequences. As the SEAM-derived entropy-based CSM captures the sequence determinants driving foreground motif activity per cluster, SEAM uses this information to separate the invariant background signal from mechanism-specific motifs (such as TF motifs) in each cluster. First, the background entropy of the sequence library is calculated using the mutation rate, *r*, used to generate the local sequence library and the corresponding probability, *p*, of a position remaining unchanged, where *p* = 1 − *r*. For a sequence with *c* nucleotides, the background entropy, *H*_BG_, is calculated as the entropy of the following distribution:

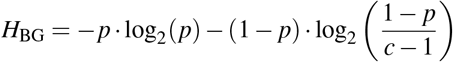

Next, an entropy threshold *H*_0_ = *H*_BG_*/*2 is set. For each averaged attribution map of a given cluster, indexed by *k*, the attribution values are set to zero for positions *i* in the associated row of the CSM where the positional sequence entropy is less than the threshold entropy, *H*_*k,i*_ *< H*_0_. Repeating this operation across all clusters and averaging the result effectively captures the attribution background, while reintegrating background attribution values that were removed from each cluster based on the presence of cluster-specific TF motifs.

Finally, the averaged attribution background is subtracted from each of the averaged attribution maps per cluster. Within a local library, the background is uniform up to a constant scaling factor. To avoid introducing excess background signal due to mismatched amplitudes, an additional per-cluster scaling factor is applied to the background before the background is subtracted. Subtraction isolates cluster-specific TF motifs by removing common background signals, thereby enhancing the specificity of the attribution maps for mechanism-specific motifs within each cluster.

### SEAM-based covariance analysis

To identify regulatory elements that covary across distinct *cis*-regulatory mechanisms, we developed a covariance-based analysis pipeline that operates on SEAM’s CSM-entropy. This method enables the discovery of regulatory elements, defined as sets of nucleotide positions whose variability patterns are coordinated across mechanism-based clusters, without prior knowledge of their position or structure.

The CSM encodes positional entropy values across sequences assigned to each mechanism-defined cluster. We compute the covariance matrix of these entropy profiles, where each entry Cov(*i, j*) quantifies how entropy at nucleotide position *i* co-varies with entropy at nucleotide position *j* across all clusters. This matrix captures positive covariances (positions that gain or lose entropy together, indicating co-regulated or modular behavior) and negative covariances (suggesting mutually exclusive regulatory features). The covariance is computed as:

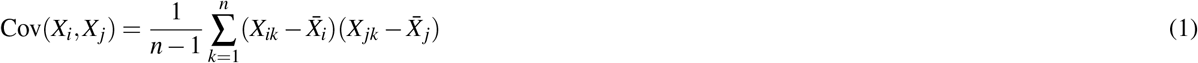

where *X*_*ik*_ denotes the entropy at position *i* in cluster *k* and 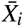 is the mean entropy at position *i* across all *K* clusters.

To isolate individual regulatory elements, we perform hierarchical clustering on the covariance matrix using average linkage. Clusters of covarying positions are extracted by cutting the dendrogram at a height that maximizes inter-cluster separation while maintaining within-element coherence. This produces a set of contiguous or semi-contiguous positional clusters, each corresponding to a putative regulatory element, whose coordinates may span flexible, motif-like regions or dispersed compositional domains.

For each regulatory element, we calculate its activity within each mechanism-defined cluster based on entropy depletion relative to background expectations. Specifically, the element’s activity in cluster *c* is defined as:

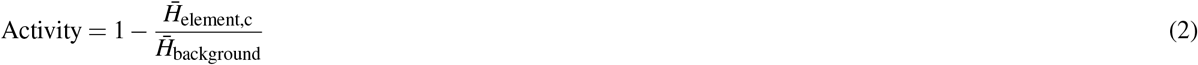

where 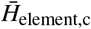 is the average entropy across the regulatory element’s positions in cluster *c*, and 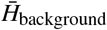 is the mean entropy of the same positions across all clusters or a null expectation under uniform mutation. This score quantifies how conserved (i.e., active) a given element is within each mechanistic context.

These activity values populate a “binding configuration matrix” in which rows represent regulatory elements defined by covariance clustering and columns correspond to SEAM-defined clusters. The matrix can be represented in binary form (active/inactive states defined by a threshold) or in continuous form to reflect graded activity. Regulatory elements are defined as sets of nucleotide positions obtained from hierarchical clustering of the covariance matrix. Each element is mapped onto attribution space by aligning its constituent positions with the corresponding attribution profiles across clusters.

### SEAM-based epistasis analysis

SEAM can quantify combinatorial interactions between regulatory elements using cluster-based predictions, each of which comprise different different motif arrangements. After elements are identified (e.g., either visually or via covariance-based analysis), each SEAM-defined cluster can be annotated by which elements are active (present) or inactive (disrupted) and used to estimate classical interaction effects grounded in functional predictions.^75^

Consider two regulatory elements, A and B (such as TF binding sites), whose presence varies in all combinations across SEAM-defined clusters. Let *y*_*S*_ denote the median DNN prediction for the cluster containing exactly the subset of elements *S* ⊆ {*A, B*}. The pairwise interaction term, which captures the non-additive interaction between A and B, is then defined using Möbius inversion:

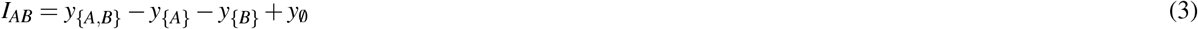

where 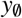 approximates the SEAM-derived background signal, based on the cluster where neither A nor B are active. A positive *I*_*AB*_ indicates synergistic behavior (greater-than-additive effects), while a negative value suggests antagonism (less-than-additive). Crucially, these mechanistic states emerge directly from SEAM’s unsupervised clustering of local sequence perturbations, bypassing the need for targeted mutagenesis or combinatorial synthesis.

This procedure extends to higher-order interactions. For three elements *A, B,C*, the third-order term is:

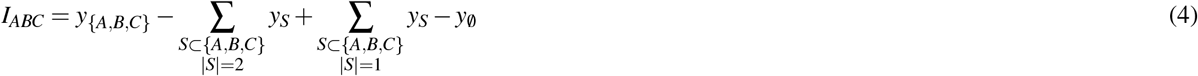

and similarly for higher cardinality, following the inclusion-exclusion principle with coefficients ( − 1)^*k*^ for subsets of size *k*. In this way, SEAM estimates epistatic contributions of regulatory elements directly from DNN predictions across clusters.

### Deep learning models

This study employed seven DNNs: DeepSTARR, CLIPNET, ProCapNet, ChromBPNet, DeepMEL2, Enformer and SpliceAI. Here, we briefly describe each DNN and how that DNN was used in our study to compute attribution maps.

- **DeepSTARR**^28^ predicts *Drosophila* enhancer activity as assayed by UMI-STARR-seq. DeepSTARR takes as input a DNA sequence of length 249 nt and outputs two scalar-valued predictions for enhancer activity for developmental (Dev) and housekeeping (Hk) regulatory programs. The DeepSTARR parameters were retrained in modern TensorFlow^76^ as specified in the original release, and the resulting model was confirmed to recapitulate the published model using performance metrics and visualization of attribution maps.
- **CLIPNET**^41^ predicts nucleotide-resolution transcription initiation profiles from a dataset consisting of matched precision run-on and 5’-capped (m^7^G) RNA enrichment (PRO-cap) and individual heterozygous human genomes from 58 genetically distinct lymphoblastoid cell lines (LCLs). CLIPNET takes as input a DNA sequence of length 1,000 nt and outputs two predictions via a “profile” head and a “counts” head. The profile head predicts strand-specific PRO-cap coverage over the central 500 nt (500 for the plus strand concatenated with 500 for the minus strand), representing the predicted base-resolution profile of initiation. The counts head predicts the total PRO-cap signal across both strands. CLIPNET is an ensemble model comprising 9 structurally identical models, each trained with a distinct holdout set of chromosomes. Unless otherwise specified, SEAM analysis was performed by averaging predictions and attribution maps across all 9 folds. Attribution analysis in CLIPNET was conducted on two-hot encoded DNA sequences, where each nucleotide at a given position is represented as a sum of two one-hot encoded nucleotides, capturing the unphased diploid sequence. When applying DeepSHAP to two-hot encoded sequences, heterozygous positions can be seen as vectors between the two orthogonal features (alleles) in the input domain. DeepSHAP evaluates the function’s behavior at this new composite point, reflecting the model’s interpretation of the combined contribution from both alleles.
- **ProCapNet**^45^ predicts nucleotide-resolution transcription initiation profiles as measured by PRO-cap in human K562 cells. ProCapNet takes as input a homozygous DNA sequence of length 2,114 nt and generates two predictions via a profile head and a counts head. The profile head predicts nucleotide-resolution initiation activity across both strands within a central 1,000 nt region, while the counts head predicts the log-transformed total number of PRO-cap reads with 5’ ends mapped within this region, summed across both strands. ProCapNet was trained using a 7-fold cross-validation scheme. Unless otherwise specified, SEAM analysis was performed by averaging predictions and attribution maps across all 7 folds. Profile head predictions were consolidated into a single explainable scalar following the approach used in the original publication.
- **ChromBPNet**^20^ predicts nucleotide-resolution chromatin accessibility profiles. ChromBPNet takes as input a DNA sequence of length 2,048 nt and generates two predictions via a “profile” head and a “counts” head. The profile head predicts nucleotide-resolution coverage within a central 2,114 nt region, while counts head predicts the natural log count of the aligned reads within this region. ChromBPNet was trained using a 5-fold cross-validation scheme. Unless otherwise specified, SEAM analysis was performed by averaging predictions and attribution maps across all 5 folds. Profile head predictions were consolidated into a single explainable scalar following the approach used in the original publication. Our primary SEAM analysis was conducted using a ChromBPNet model trained on DNase-seq in THP-1 cells, available at 10.5281/zenodo.10403551. To assess the generality of SEAM background signals, we additionally applied SEAM to alternative versions of ChromBPNet trained on: (1) DNase-seq in K562 cells, (2) ATAC-seq in K562 cells, and (3) DNase-seq in THP-1 cells using the chrombpnet_nobias model trained to predict bias corrected accessibility profiles. The models trained on ATAC-seq and DNase-seq profiles from K562 cells are available at the ENCODE portal: http://encodeproject.org, under file IDs ENCFF984RAF and ENCFF574YLK, respectively.
- **DeepMEL2**^49^ predicts melanoma-specific chromatin accessibility as measured by ATAC-seq and allele-specific chromatin accessibility variants. DeepMEL2 takes as input the forward and reverse DNA strands, each of length 500 nt, and outputs a vector of binarized predictions across 47 classes, each representing a melanoma *cis*-regulatory topic. Only class 16 (MEL) was used in this work. Contribution scores were generated for each strand separately, and following previous work,^49^ we averaged the contribution scores over both strands to visualize attribution maps.
- **Enformer**^18^ predicts many different types of functional genomic track (for example, ChIP–seq, DNase-seq, and ATAC–seq) across the human and mouse genomes. Enformer takes as input a DNA sequence of length 393,216 nt and (for humans) outputs 5,313 profiles (one for each track) where each profile comprises 128 bins, each bin spanning 32 nt, representing the central 114,688 nt of the input sequence. The published Enformer parameters were used to compute these profiles. For the CSM shown in Supplementary Fig. 1e, we used human predictions, where saliency maps were computed as in the original study by selecting all CD14-positive DNase tracks and creating a target mask for a 16-bin (2,048 nt) region centered at the enhancer, then computing gradients fo the summed predictions with respect to the input sequence.
- **SpliceAI**^77^ predicts splice donor and acceptor sites directly from primary pre-mRNA sequences using a deep residual neural network. The model takes as input a sequence of 10,000 nucleotides flanking each candidate site (i.e., ±5 kb context) and produces position-wise scores indicating the likelihood of splice acceptor or donor activity. For the CSM shown in Supplementary Fig. 1b, saliency maps were computed using gradient-based attribution by selecting the maximum prediction score across all positions for the acceptor site class, then computing gradients of this maximum prediction with respect to the input sequence, with the resulting attribution maps averaged across 5 ensemble models.

### SEAM API

SEAM takes as input a reference sequence, a sequence-function model (oracle), an attribution method, and a clustering strategy. The SEAM API provides optimized GPU and CPU support across six modular components:

- **Mutagenizer:** Applies a user-specified mutagenesis strategy to generate an *in silico* sequence library consisting of *N* aligned sequence variants derived from the reference sequence. This class is imported from the SQUID library,^32^ and supports generation of different libraries, including local, global, optimized, and complete. The complete library also supports all combinatorial mutations up to a specified mutation order (e.g., all single- and double-nucleotide variants). GPU-acceleration and batch processing are supported for all operations, along with optimized CPU support. Additional details can be found in the section “SEAM libraries”.
- **Compiler:** Standardizes sequence analysis data by converting one-hot encoded sequences to string format and computing associated metrics. Compiles sequence and functional properties into a DataFrame, with options to include calculations for Hamming distances relative to the reference sequence and global importance analysis^78^ (GIA) scores computed with respect to background predictions. The implementation includes GPU-accelerated sequence conversion and vectorized operations for efficient processing of large datasets.
- **Attributer:** Computes attribution maps for each of the *N* sequences using the specified attribution method, including Saliency Maps, Integrated Gradients (IntGrad), SmoothGrad, DeepSHAP, and *in silico Mutagenesis* (ISM). Each attribution map quantifies the base-wise contribution to the model’s predicted regulatory activity. Algorithms for ISM, Saliency Maps, SmoothGrad, and Integrated Gradients were refactored for TensorFlow 2^76^ within the SEAM API as class-based objects optimized for GPU-accelerated batch processing. DeepSHAP was integrated as a class-based object from the publicly available kundajelab-shap repository, which lacks GPU-accelerated batch processing over inputs. For PyTorch^79^ models such as ProCapNet,^45^ we applied DeepSHAP using the DeepLiftShap implementation provided in Captum.^80^ Efficient batch processing and flexible sequence windowing is also supported for analyzing large datasets. For baseline-specific attribution methods, multiple baseline types are available (zero baseline, random shuffle, or dinucleotide-preserved shuffle) to account for different biological assumptions about the reference state. Additional details can be found in the section “Attribution methods”.
- **Clusterer:** Clusters the resulting attribution maps to identify qualitatively distinct regulatory mechanisms. Maps can optionally be embedded into a low-dimensional space prior to clustering to extend interpretability or improve scalability. The class supports multiple embedding methods including UMAP, t-SNE, and PCA, with GPU acceleration available for computationally intensive operations. Clustering can be performed using hierarchical clustering (with GPU optimization), K-means, or DBSCAN algorithms. The implementation includes memory-efficient batch processing for large datasets and provides comprehensive visualization tools for analyzing clustering results, including embedding plots, 2D histograms, and dendrograms. The class automatically handles dependency management and falls back to CPU implementations when GPU libraries are unavailable. Additional details can be found in the section “Clustering methods”.
- **MetaExplainer:** Averages attribution maps within each cluster to generate cluster-averaged attribution maps, yielding noise-reduced representations of distinct regulatory mechanisms. Sequence cluster assignments are then used to compute position-wise nucleotide frequencies for each cluster, together forming the Cluster Summary Matrix (CSM), which can be computed using either positional Shannon entropy or percent mismatch relative to the reference. The class implements background separation to remove non-specific signal patterns, with optional adaptive scaling to account for cluster-specific background magnitudes. Comprehensive visualization tools are also provided, including sequence logos, attribution logos, and cluster statistics plots, with support for both PWM-based and enrichment-based sequence analysis. The implementation includes GPU acceleration and batch processing for computationally intensive operations, with fallbacks to optimized CPU support. Additional details can be found in the section “SEAM-based background separation”.
- **Identifier:** Uses the cluster-averaged attribution maps and CSM to identify motifs, delimit their positions, and separate foreground (mutation-sensitive) from background (mutation-insensitive) signals using background subtraction. The class implements hierarchical clustering of position-wise covariance patterns to identify distinct regulatory elements and their positions. It provides comprehensive visualization tools including covariance matrices, dendrograms, and binding configuration matrices showing TFBS activity across clusters. The implementation supports both binary and continuous activity modes. The class includes memory-efficient matrix operations and flexible visualization options with support for view windows and customizable styling. Additional details can be found in the section “SEAM-based covariance analysis”.

Comprehensive documentation for all classes, including detailed API references, usage examples, and tutorials, is available at https://seam-nn.readthedocs.io/.

### SEAM GUI

To facilitate exploration and interpretation of SEAM outputs, we developed the SEAM Interactive Interpretability Tool, a graphical user interface (GUI) that enables intuitive navigation and analysis of mechanisms in attribution space (Supplementary Fig. 21). The tool accepts SEAM’s core outputs: a sequence library, attribution maps, and either a linkage matrix or an embedded representation of attribution space (e.g., UMAP or t-SNE).

If an embedding is provided, the GUI renders this attribution space, in which each point represents an attribution map derived from a sequence variant. Users may overlay clustering labels (e.g., from K-means or DBSCAN), or draw custom boundaries directly within the interface. For each selected cluster, the GUI displays averaged attribution maps, predicted activity distributions, and associated summary statistics.

Points in the attribution space can be dynamically colored by predicted activity, Hamming distance to the reference, or a 2D occupancy histogram that highlights the density of mechanisms in attribution space. These visualizations enable users to interpret spatial relationships between mechanism and function, detect sparsely or densely sampled mechanisms, and identify mutation-driven transitions.

The GUI also includes an interactive Cluster Summary Matrix viewer, available for both linkage- and embedding-based clustering strategies. Each CSM cell can be clicked to display the full nucleotide distribution at that position within the selected cluster, with annotations relative to the reference sequence. This functionality helps uncover mutation hotspots, motif architecture, and the sequence features responsible for mechanistic divergence.

Together, these tools provide an accessible and hypothesis-driven interface for interpreting the full complexity of SEAM’s output, bridging automated clustering with exploratory analysis of regulatory mechanism diversity.

## Data availability

The datasets used in this study are publicly available from established sources. PBM data were obtained from the UniPROBE database.^81^ Genomic sequences are based on the hg38 human reference genome. Additional sequence data, including test sets held out during the training of models such as DeepSTARR, were accessed directly from the respective model repositories.

## Code availability

SEAM is an open-source Python package based on TensorFlow,^76^ and contains CPU and GPU-optimized code for attribution analysis and clustering. SEAM can be installed via pip (https://pypi.org/project/seam-nn) or GitHub (https://github.com/evanseitz/sean-nn). The GitHub repository contains links to running several examples from our analysis in Google Colab. Documentation is provided on ReadTheDocs (https://seam-nn.readthedocs.io).

## Acknowledgements

We thank Charles Danko, Adam He, Jack Desmarais, Kai Loell, Zhihan Liu, Julia Zeitlinger and Jesse Engreitz for helpful discussions. This work was supported in part by: the Simons Center for Quantitative Biology at Cold Spring Harbor Laboratory; NIH grants R01HG012131 (P.K.K, E.E.S., J.B.K., D.M.M.), R01HG011787 (J.B.K., E.E.S., D.M.M.), R01GM149921 (P.K.K.), R35GM133777 (J.B.K.), F32HG013265 (E.E.S.), and R35GM133613 (D.M.M.). CSHL Cancer Center Developmental Funds partly supported this work from the Cancer Center Support Grant P30CA045508, and computations were performed using equipment supported by NIH grant S10OD028632.

## Author contributions

E.E.S. conceived the method, developed the software, and performed the analysis. E.E.S., D.M.M., J.B.K., and P.K.K. designed the study and wrote the paper. J.B.K. and P.K.K. supervised the research.

## Competing interests

The authors declare no competing interests.

## Notes

### Competing Interest Statement

The authors have declared no competing interest.

## References

1. Banerji, J., Rusconi, S. & Schaffner, W. Expression of a beta-globin gene is enhanced by remote SV40 DNA sequences. Cell 27, 299–308, DOI: 10.1016/0092-8674(81)90413-X (1981). Doi: 10.1016/0092-8674(81)90413-X.

2. Spitz, F. & Furlong, E. E. M. Transcription factors: From enhancer binding to developmental control. Nat. Rev. Genet. 13, 613–626, DOI: 10.1038/nrg3207 (2012).

3. Rickels, R. & Shilatifard, A. Enhancer logic and mechanics in development and disease. Trends Cell Biol. 28, 608–630, DOI: 10.1016/j.tcb.2018.04.003 (2018).

4. Shlyueva, D., Stampfel, G. & Stark, A. Transcriptional enhancers: From properties to genome-wide predictions. Nat. Rev. Genet. 15, 272–286, DOI: 10.1038/nrg3682 (2014).

5. Nagy, G. & Nagy, L. Motif grammar: The basis of the language of gene expression. Comput. Struct. Biotechnol. J. 18, 2026–2032, DOI: 10.1016/j.csbj.2020.07.007 (2020). ECollection 2020.

6. Zeitlinger, J. Seven myths of how transcription factors read the cis-regulatory code. Curr. opinion systems biology 23, 22–31 (2020).

7. Mariani, L. et al. DNA bendability regulates transcription factor binding to nucleosomes. Nat. Struct. & Mol. Biol. 1–11 (2025).

8. Kim, S. & Wysocka, J. Deciphering the multi-scale, quantitative cis-regulatory code. Mol. Cell 83, 373–392, DOI: 10.1016/j.molcel.2022.12.032 (2023). Reimagining the Central Dogma.

9. Fuqua, T. et al. Dense and pleiotropic regulatory information in a developmental enhancer. Nature 587, 235–239, DOI: 10.1038/s41586-020-2816-5 (2020). Epub 2020 Oct 14.

10. Martyn, G. E. et al. Rewriting regulatory DNA to dissect and reprogram gene expression. Cell 0, —, DOI: 10.1016/j.cell.2025.04.016 (2025). Online ahead of print.

11. Crutchfield, J. P. & van Nimwegen, E. The evolutionary unfolding of complexity. In Landweber, L. F. & Winfree, E. (eds.) Evolution as Computation, 67–94 (Springer Berlin Heidelberg, Berlin, Heidelberg, 2002).

12. Levine, M. Transcriptional enhancers in animal development and evolution. Curr. Biol. 20, R754–R763, DOI: 10.1016/j.cub.2010.06.070 (2010). Doi: 10.1016/j.cub.2010.06.070.

13. Franchini, L. F. & Pollard, K. S. Human evolution: The non-coding revolution. BMC biology 15, 1–12 (2017).

14. Claussnitzer, M. et al. A brief history of human disease genetics. Nature 577, 179–189, DOI: 10.1038/s41586-019-1879-7 (2020).

15. Nasser, J. et al. Genome-wide enhancer maps link risk variants to disease genes. Nature 593, 238–243, DOI: 10.1038/s41586-021-03446-x (2021).

16. Maurano, M. T. et al. Systematic localization of common disease-associated variation in regulatory DNA. Science 337, 1190–1195, DOI: 10.1126/science.1222794 (2012). https://www.science.org/doi/pdf/10.1126/science.1222794.

17. Horn, S. et al. TERT promoter mutations in familial and sporadic Melanoma. Science 339, 959–961, DOI: 10.1126/science.1230062 (2013).

18. Avsec, Ž, et al. Effective gene expression prediction from sequence by integrating long-range interactions. Nat. Methods 18, 1196–1203, DOI: 10.1038/s41592-021-01252-x (2021).

19. Toneyan, S., Tang, Z. & Koo, P. K. Evaluating deep learning for predicting epigenomic profiles. Nat. Mach. Intell. 4, 1088–1100 (2022).

20. Pampari, A. et al. ChromBPNet: bias factorized, base-resolution deep learning models of chromatin accessibility reveal cis-regulatory sequence syntax, transcription factor footprints and regulatory variants. BioRxiv 2024–12 (2025).

21. Jaganathan, K. et al. Predicting expression-altering promoter mutations with deep learning. Science eads7373 (2025).

22. Avsec, Ž. et al. AlphaGenome: advancing regulatory variant effect prediction with a unified DNA sequence model. bioRxiv 2025–06 (2025).

23. Zhou, J. & Troyanskaya, O. G. Predicting effects of noncoding variants with deep learning-based sequence model. Nat. Methods 12, 931–934 (2015).

24. Simonyan, K., Vedaldi, A. & Zisserman, A. Deep inside convolutional networks: Visualising image classification models and saliency maps. In Workshop at International Conference on Learning Representations (2014).

25. Lundberg, S. M. & Lee, S.-I. A unified approach to interpreting model predictions. In Advances in Neural Information Processing Systems, vol. 30 (Curran Associates, Inc., 2017).

26. Koo, P. K. & Ploenzke, M. Deep learning for inferring transcription factor binding sites. Curr. Opin. Syst. Biol. 19, 16–23 (2020).

27. Novakovsky, G., Dexter, N., Libbrecht, M. W., Wasserman, W. W. & Mostafavi, S. Obtaining genetics insights from deep learning via explainable artificial intelligence. Nat. Rev. Genet. 24, 125–137 (2022).

28. de Almeida, B. P., Reiter, F., Pagani, M. & Stark, A. DeepSTARR predicts enhancer activity from DNA sequence and enables the de novo design of synthetic enhancers. Nat. Genet. 54, 613–624 (2022).

29. Shrikumar, A. et al. Technical note on transcription factor motif discovery from importance scores (TF-MoDISco) version 0.5.6.5 (2020). 1811.00416.

30. Avsec, Ž. et al. Base-resolution models of transcription-factor binding reveal soft motif syntax. Nat. Genet. 53, 354–366 (2021).

31. Horton, C. A. et al. Short tandem repeats bind transcription factors to tune eukaryotic gene expression. Science 381, eadd1250 (2023).

32. Seitz, E., McCandlish, D., Kinney, J. & Koo, P. Interpreting cis-regulatory mechanisms from genomic deep neural networks using surrogate models. Nat. Mach. Intell. DOI: 10.1038/s42256-024-00851-5 (2024).

33. Dalal, K. et al. Interpreting regulatory mechanisms of Hippo signaling through a deep learning sequence model. Cell Genomics (2025).

34. Jr., J. H. W. Hierarchical grouping to optimize an objective function. J. Am. Stat. Assoc. 58, 236–244, DOI: 10.1080/01621459.1963.10500845 (1963).

35. de Almeida, B. P. et al. Targeted design of synthetic enhancers for selected tissues in the Drosophila embryo. Nature 626, 207–211, DOI: 10.1038/s41586-023-06905-9 (2024).

36. Reiter, F., de Almeida, B. P. & Stark, A. Enhancers display constrained sequence flexibility and context-specific modulation of motif function. Genome Res. 33, 346–358, DOI: 10.1101/gr.277246.122 (2023). http://genome.cshlp.org/content/33/3/346.full.pdf+html.

37. Kudla, G., Lipinski, L., Caffin, F., Helwak, A. & Zylicz, M. High guanine and cytosine content increases mRNA levels in mammalian cells. PLOS Biol. 4, null, DOI: 10.1371/journal.pbio.0040180 (2006).

38. Tillo, D. & Hughes, T. R. G+C content dominates intrinsic nucleosome occupancy. BMC Bioinforma. 10, 442, DOI: 10.1186/1471-2105-10-442 (2009).

39. Fenouil, R. et al. CpG islands and GC content dictate nucleosome depletion in a transcription-independent manner at mammalian promoters. Genome Res. 22, 2399–2408, DOI: 10.1101/gr.138776.112 (2012).

40. Castellanos, M., Mothi, N. & Muñoz, V. Eukaryotic transcription factors can track and control their target genes using DNA antennas. Nat. Commun. 11, 540, DOI: 10.1038/s41467-019-14217-8 (2020).

41. He, A. Y. & Danko, C. G. Dissection of core promoter syntax through single nucleotide resolution modeling of transcription initiation. bioRxiv DOI: 10.1101/2024.03.13.583868 (2024).

42. Gupta, H. et al. Highly diversified core promoters in the human genome and their effects on gene expression and disease predisposition. BMC Genomics 21, 842, DOI: 10.1186/s12864-020-07222-5 (2020).

43. Zhou, J. et al. Genetic and bioinformatic analyses of the expression and function of PI3K regulatory subunit PIK3R3 in an Asian patient gastric cancer library. BMC Med. Genomics 5, 34, DOI: 10.1186/1755-8794-5-34 (2012).

44. Wang, G. et al. PIK3R3 induces epithelial-to-mesenchymal transition and promotes metastasis in colorectal cancer. Mol. Cancer Ther. 13, 1837–1847, DOI: 10.1158/1535-7163.MCT-14-0049 (2014). https://aacrjournals.org/mct/article-pdf/13/7/1837/2329174/1837.pdf.

45. Cochran, K. et al. Dissecting the cis-regulatory syntax of transcription initiation with deep learning. bioRxiv DOI: 10.1101/2024.05.28.596138 (2024).

46. Badis, G. et al. Diversity and complexity in DNA recognition by transcription factors. Science 324, 1720–1723, DOI: 10.1126/science.1162327 (2009).

47. de Boer, C. G. et al. Deciphering eukaryotic gene-regulatory logic with 100 million random promoters. Nat. Biotechnol. 38, 56–65, DOI: 10.1038/s41587-019-0315-8 (2020).

48. Gosai, S. et al. Machine-guided design of synthetic cell type-specific cis-regulatory elements. bioRxiv DOI: 10.1101/2023.08.08.552077 (2023).

49. Taskiran, I. I. et al. Cell-type-directed design of synthetic enhancers. Nature 626, 212–220, DOI: 10.1038/s41586-023-06936-2 (2024).

50. Schreiber, J., Lu, Y. Y. & Noble, W. S. Ledidi: Designing genomic edits that induce functional activity. bioRxiv DOI: 10.1101/2020.05.21.109686 (2020). https://www.biorxiv.org/content/early/2020/05/25/2020.05.21.109686.full.pdf.

51. Atak, Z. K. et al. Interpretation of allele-specific chromatin accessibility using cell state-aware deep learning. Genome Res. 31, 1082–1096, DOI: 10.1101/gr.260851.120 (2021).

52. Tuğrul, M., Paixao, T., Barton, N. H. & Tkačik, G. Dynamics of transcription factor binding site evolution. PLoS genetics 11, e1005639 (2015).

53. Mathelier, A. et al. DNA shape features improve transcription factor binding site predictions in vivo. Cell Syst. 3, 278–286.e4, DOI: 10.1016/j.cels.2016.07.001 (2016).

54. Sielemann, J., Wulf, D., Schmidt, R. & Bräutigam, A. Local DNA shape is a general principle of transcription factor binding specificity in Arabidopsis thaliana. Nat. Commun. 12, 6549, DOI: 10.1038/s41467-021-26819-2 (2021).

55. Ghoshdastidar, D. & Bansal, M. Flexibility of flanking DNA is a key determinant of transcription factor affinity for the core motif. Biophys. J. 121, 3987–4000, DOI: 10.1016/j.bpj.2022.08.015 (2022).

56. O’Dwyer, M. R. et al. Nucleosome fibre topology guides transcription factor binding to enhancers. Nature 638, 251–260, DOI: 10.1038/s41586-024-08333-9 (2025).

57. Toneyan, S. & Koo, P. K. Interpreting cis-regulatory interactions from large-scale deep neural networks. Nat. genetics 56, 2517–2527 (2024).

58. Koo, P. K., Anand, P., Paul, S. B. & Eddy, S. R. Inferring sequence-structure preferences of rna-binding proteins with convolutional residual networks. BioRxiv 418459 (2018).

59. Tomaz da Silva, P. et al. Nucleotide dependency analysis of dna language models reveals genomic functional elements. bioRxiv 2024–07 (2024).

60. Janizek, J. D., Sturmfels, P. & Lee, S.-I. Explaining explanations: Axiomatic feature interactions for deep networks. J. Mach. Learn. Res. 22, 1–54 (2021).

61. Marshall, D. et al. The structure-fitness landscape of pairwise relations in generative sequence models. BioRxiv 2020–11 (2020).

62. Zhang, Z. et al. Protein language models learn evolutionary statistics of interacting sequence motifs. Proc. Natl. Acad. Sci. 121, e2406285121 (2024).

63. Smilkov, D., Thorat, N., Kim, B., Viégas, F. & Wattenberg, M. SmoothGrad: Removing noise by adding noise. arXiv (2017).

64. Sundararajan, M., Taly, A. & Yan, Q. Axiomatic attribution for deep networks. arXiv (2017).

65. Shrikumar, A., Greenside, P. & Kundaje, A. Learning important features through propagating activation differences. In International Conference on Machine Learning, 3145–3153 (PMlR, 2017).

66. Majdandzic, A., Rajesh, C. & Koo, P. K. Correcting gradient-based interpretations of deep neural networks for genomics. Genome Biol. 24, 109 (2023).

67. Posfai, A., Zhou, J., McCandlish, D. M. & Kinney, J. B. Gauge fixing for sequence-function relationships. PLoS Comput. Biol. 21, e1012818 (2025).

68. Virtanen, P. et al. SciPy 1.0: Fundamental Algorithms for Scientific Computing in Python. Nat. Methods 17, 261–272, DOI: 10.1038/s41592-019-0686-2 (2020).

69. Lloyd, S. Least squares quantization in PCM. IEEE Transactions on Inf. Theory 28, 129–137, DOI: 10.1109/TIT.1982.1056489 (1982).

70. Ester, M., Kriegel, H.-P., Sander, J., Xu, X. et al. A density-based algorithm for discovering clusters in large spatial databases with noise. In kdd, vol. 96, 226–231 (1996).

71. Pearson, K. LIII. On lines and planes of closest fit to systems of points in space. The London, Edinburgh, Dublin Philos. Mag. J. Sci. 2, 559–572, DOI: 10.1080/14786440109462720 (1901).

72. van der Maaten, L. & Hinton, G. Visualizing data using t-SNE. J. Mach. Learn. Res. 9, 2579–2605 (2008).

73. McInnes, L., Healy, J. & Melville, J. UMAP: Uniform manifold approximation and projection for dimension reduction (2020). 1802.03426.

74. Buitinck, L. et al. API design for machine learning software: Experiences from the scikit-learn project. In ECML PKDD Workshop: Languages for Data Mining and Machine Learning, 108–122 (2013).

75. Phillips, P. C. Epistasis — the essential role of gene interactions in the structure and evolution of genetic systems. Nat. Rev. Genet. 9, 855–867, DOI: 10.1038/nrg2452 (2008).

76. Abadi, M. et al. TensorFlow: Large-scale machine learning on heterogeneous systems (2015). Software available from tensorflow.org.

77. Jaganathan, K. et al. Predicting splicing from primary sequence with deep learning. Cell 176, 535–548.e24, DOI: 10.1016/j.cell.2018.12.015 (2019).

78. Koo, P. K., Majdandzic, A., Ploenzke, M., Anand, P. & Paul, S. B. Global importance analysis: An interpretability method to quantify importance of genomic features in deep neural networks. PLoS Comput. Biol. 17, e1008925 (2021).

79. Paszke, A. et al. Automatic differentiation in PyTorch. (2017).

80. Kokhlikyan, N. et al. Captum: A unified and generic model interpretability library for PyTorch (2020). Cite arxiv:2009.07896.

81. Newburger, D. E. & Bulyk, M. L. UniPROBE: an online database of protein binding microarray data on protein–DNA interactions. Nucleic Acids Res. 37, D77–D82, DOI: 10.1093/nar/gkn660 (2008).

