## Supplemental Information for "Uncovering the Mechanistic Landscape of Regulatory DNA with Deep Learning"

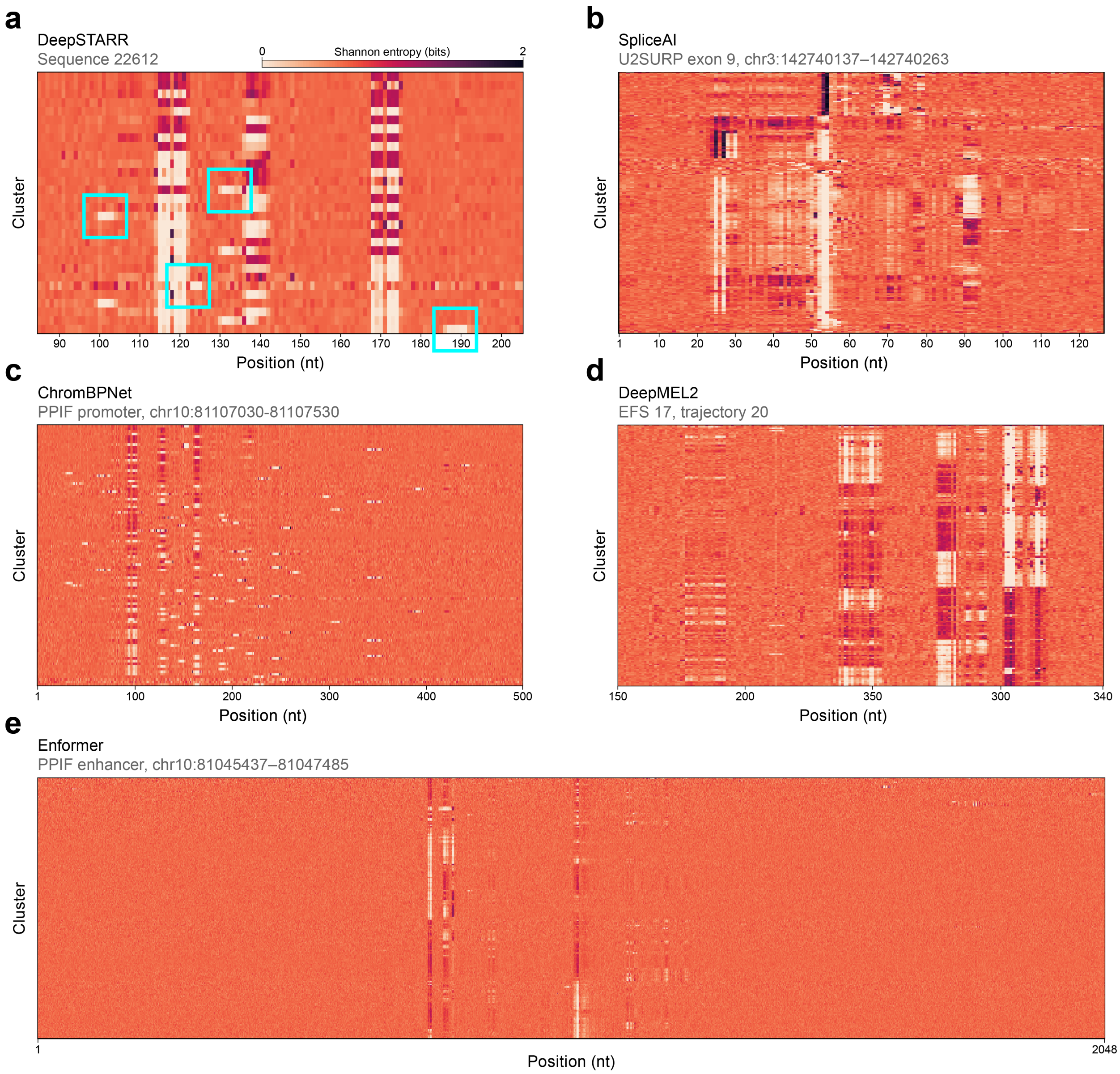


**Supplementary Figure 1. SEAM entropy-based CSM patterns are consistent across genomic DNNs, data modalities, and species.** SEAM entropy-based cluster summary matrices (CSM) generated with SEAM using different genomic DNNs, with each CSM computed based on a local library (10% mutation rate) corresponding to a unique reference sequence. **a**, CSM for the *Drosophila* enhancer shown in Fig. 2b, generated using the DeepSTARR model with DeepSHAP attribution maps (using the developmental head) and hierarchical clustering. Cyan rectangles highlight examples of motif-preserving signals that appear outside of vertical bands, each corresponding to the emergence of a *de novo* motif for the respective cluster(s). **b**, CSM for the human U2SURP exon 9, generated using the SpliceAI model with Saliency attribution maps (using the max prediction at the acceptor head; see Methods) and K-Means clustering on the UMAP embedding. **c**, CSM for the human PPIF promoter, generated using the ChromBPNet model with DeepSHAP attribution maps (using the counts head) and hierarchical clustering. **d**, CSM for the final trajectory from the Evolved from Scratch (EFS) sequence 17, generated using the DeepMEL2 model with DeepSHAP attribution maps (using class 16) and K-Means clustering on the UMAP embedding. **e**, CSM for the human PPIF enhancer, generated using the Enformer model with Saliency attribution maps (using all human DNase CD14-positive profiles; see Methods) and K-Means clustering on the UMAP embedding. CSM clusters for DeepSTARR and ChromBPNet examples are ordered by ascending DNN prediction. CSM clusters for SpliceAI, DeepMEL2, and Enformer examples are ordered according to entropy-similarity over positions (using hierarchical clustering) to better visualize entropy-based patterns.

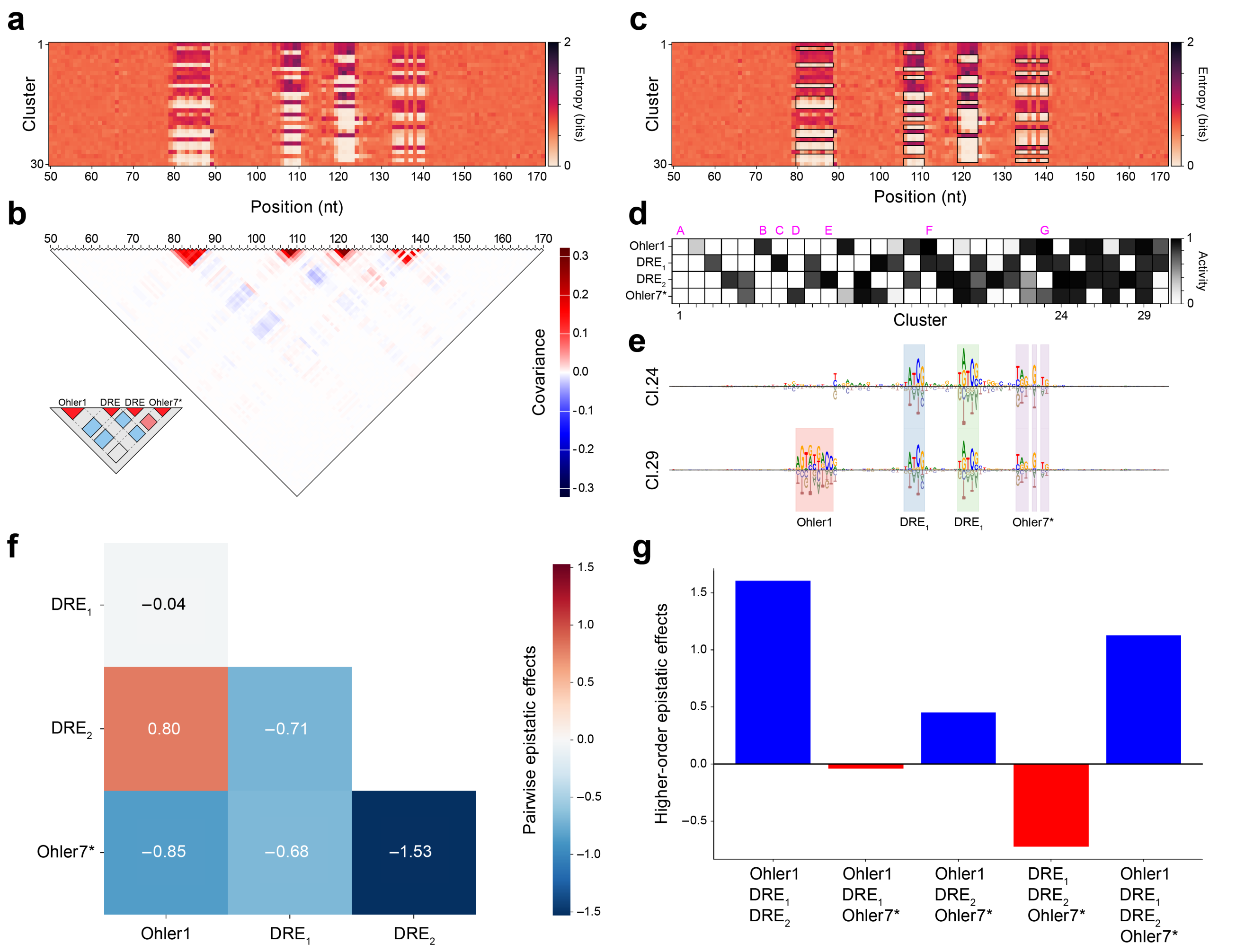


**Supplementary Figure 2. SEAM covariance analysis annotates mutation-sensitive regulatory elements and quantifies combinatorial logic.** **a**, SEAM entropy-based CSM for the DeepSTARR locus shown in Fig. 2a. **b**, Covariance matrix of the CSM, capturing how nucleotide positions co-vary in their entropy profiles across clusters (see "SEAM-based covariance analysis" in Methods). For covariance analysis, SEAM applies a second, independent hierarchical clustering to the covariance matrix (here cut into four groups corresponding to four regulatory motifs) to group together nucleotide positions that respond jointly to sequence perturbation. Inset: covariance patterns also reveal co-occurrence trends between regulatory motifs across mechanistically-defined clusters. **c**, CSM with low-entropy regions outlined in black, with the horizontal extent of each region determined by the nucleotide groups from the covariance-based clustering, and the vertical boundaries defined by an entropy threshold. **d**, Low-entropy regions within each black rectangle identified in the CSM are averaged and transformed into an activity level for each TFBS, forming a binding configuration matrix. Magenta letters at the top of the binding configuration matrix denote the position of example clusters in Fig. 2a. Some states appear as graded versions of others (e.g., Ohler1 in cluster 2 vs. cluster 6), reflecting contextual modulation of attribution amplitudes. **e**, The position of regulatory elements in each state are combined with their activity in that state to highlight active motifs in the cluster-averaged attribution maps. Attribution maps are cluster-averaged and background separated (see Methods). **f**, Pairwise epistatic interaction matrix computed using Möbius inversion (see "SEAM-based epistasis analysis" in Methods), using median DNN predictions corresponding to each optimal match in the binding configuration matrix. **g**, Bar plot of higher-order epistatic effects for all combinations of regulatory elements.

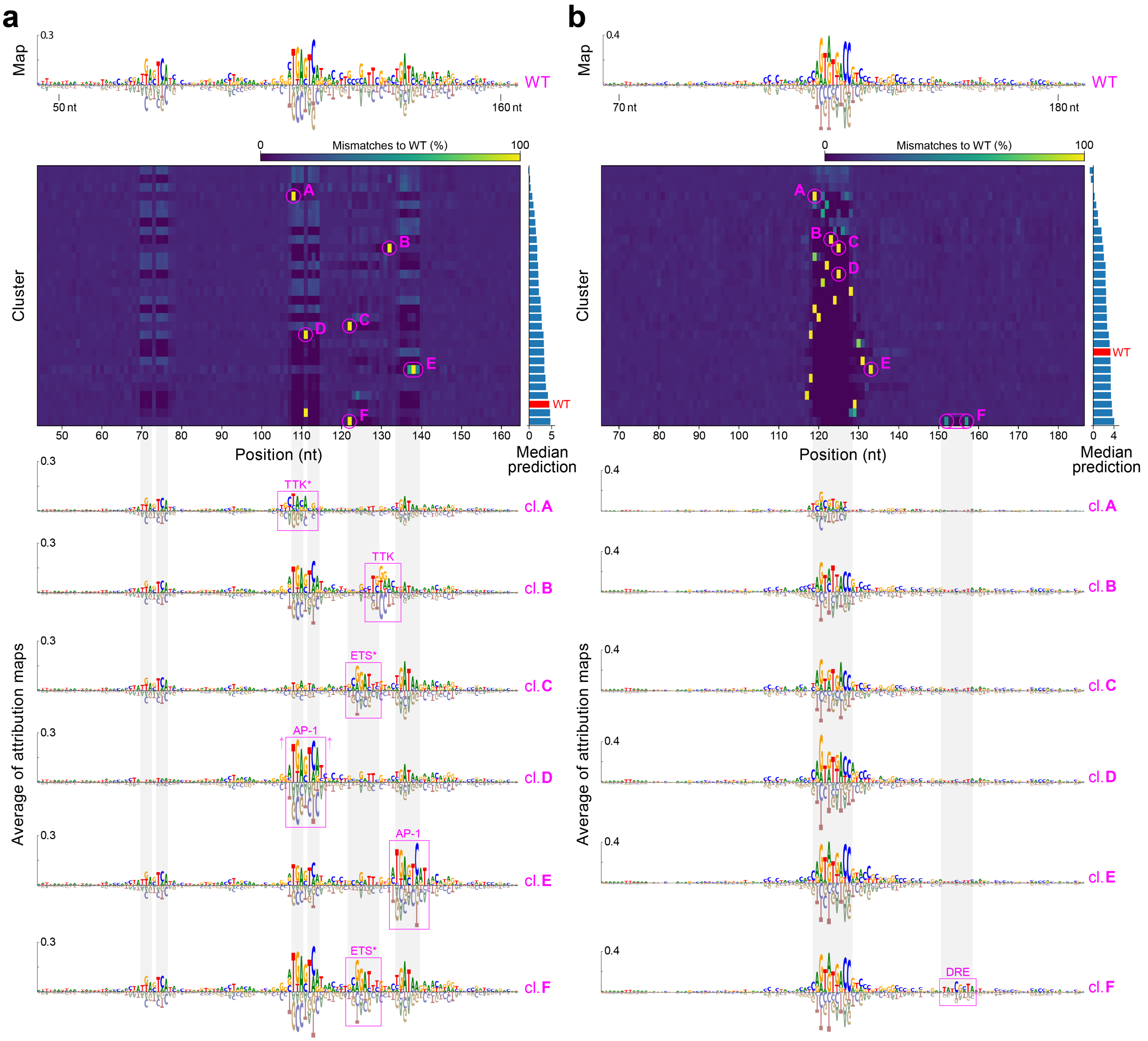


**Supplementary Figure 3. SEAM identifies driver mutations that reprogram regulatory mechanisms. a**, Top: CSM based on percent mismatches to the wild-type (WT) sequence for a locus from the DeepSTARR test set (index 21069), using the developmental head. Specific single-nucleotide variants (SNVs) in the CSM (middle) are associated with distinct shifts away from the WT mechanism—visualized by the average of attribution maps in each cluster (bottom)—often through the creation of poised motifs that modulate enhancer activity. For example, SNVs in clusters A and B reveal alternative binding sites for the repressor TTK that strongly or moderately repress activity, respectively. Not all driver mutations form *de novo* motifs; in cluster D, a substitution at the central position of the AP-1 TFBS increases the overall importance of all nucleotides in the core motif. **b**, CSM for a second locus from the DeepSTARR test set (index 22627), analyzed with the housekeeping head. Cluster F reveals a poised DRE TFBS governed by two driver mutations that are associated with an increase in predicted activity, while clusters A–E demonstrate fine-scale shifts in the conformation of a central Ohler1 motif, each driven by a distinct SNV.

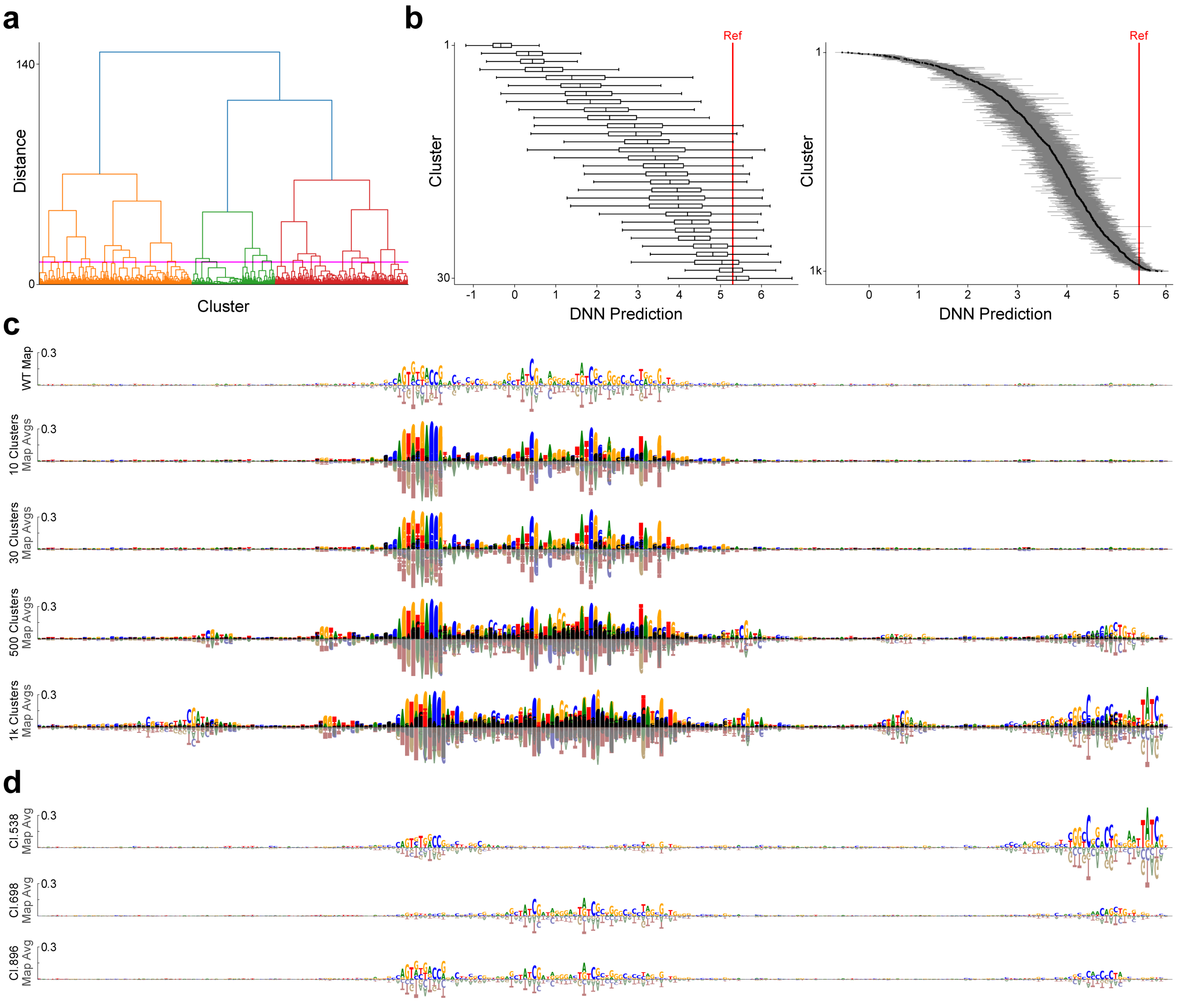


**Supplementary Figure 4. Impact of maximum cluster number on SEAM mechanistic insights. a**, SEAM dendrogram for the DeepSTARR locus shown in Fig. 2a, generated using hierarchical clustering with Ward's linkage on a Euclidean distance matrix. The pink horizontal line indicates the cut level for selecting the 30 highest-level clusters. **b**, Median DNN predictions for the 30 (left) and 1000 (right) highest-level clusters. Both plots demonstrate that the clusters span a dynamic range of DNN predictions, with finer granularity as the number of clusters increases. Lines represent the upper and lower quartiles. **c**, Comparison of the wild-type (WT) map to the overlay of all average (avg.) maps from each cluster, as the maximum number of clusters increases from 10 to 1000. As seen with 500 and 1000 clusters, more mechanisms are identified. **d**, Examples of individual mechanisms obtained using 1000 clusters, revealing new instances of TFBSs not observed at the other cut levels shown.

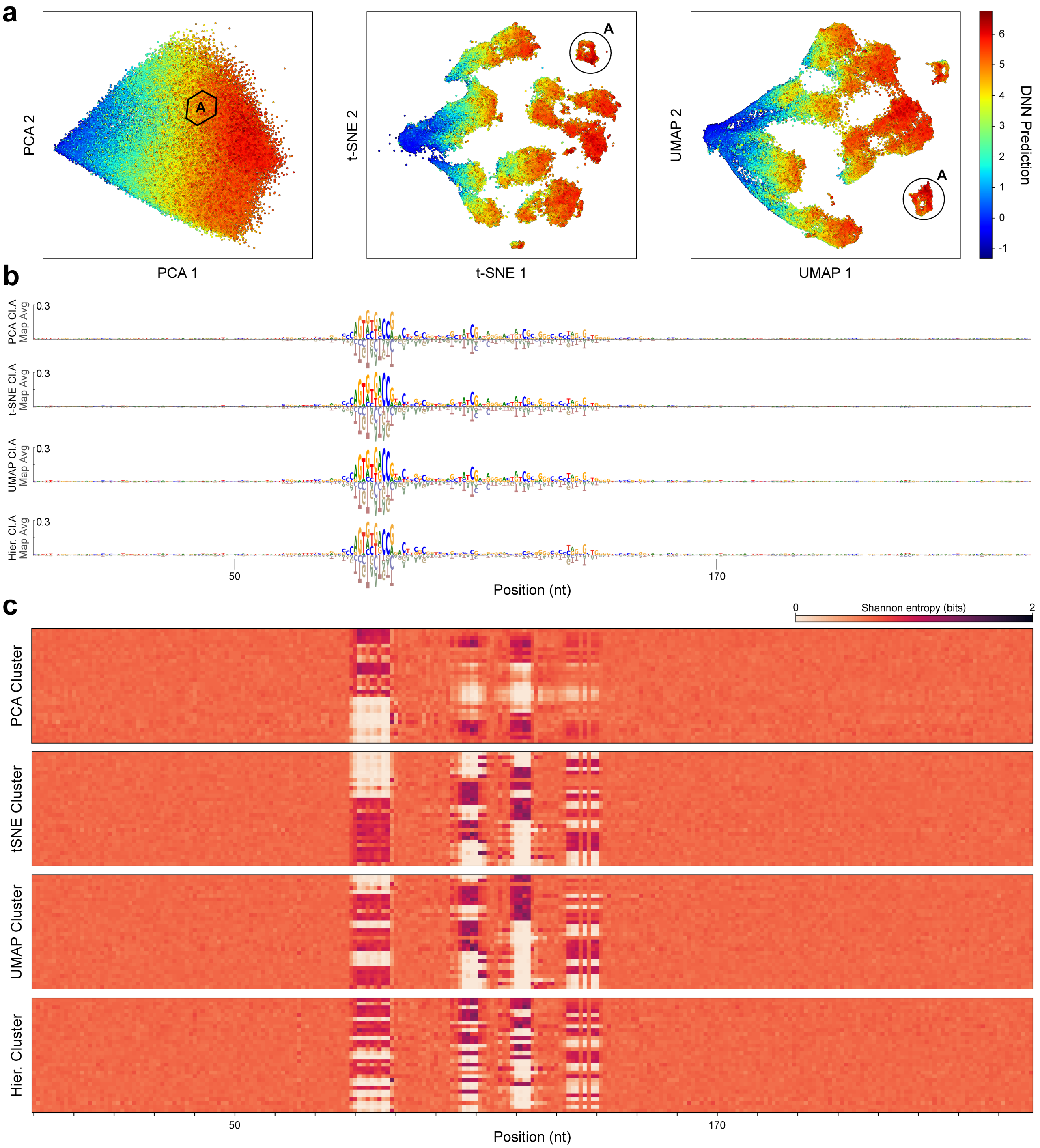


**Supplementary Figure 5. Impact of clustering method on SEAM mechanistic insights. a**, Comparison of SEAM embeddings of attribution maps for the DeepSTARR locus shown in Fig. 2a, generated using PCA, t-SNE, and UMAP. For each embedding, K-means was performed with 200 clusters. For each set of clusters in each embedding, the position of an example cluster (cl.A) is encircled, corresponding to a mechanism with a highly similar visual appearance across all three embeddings. **b**, Sequence logo for the mechanism corresponding to cluster A, generated by averaging the attribution maps in cluster A for each embedding (map avg.). **c**, Comparison of entropy-based CSMs for the three embeddings. The CSM created using hierarchical clustering, as shown in Fig. 2a, is also shown for comparison. By visual inspection, PCA with K-means produces the CSM with the lowest-resolution features, while hierarchical clustering produces the CSM with the highest-resolution features.

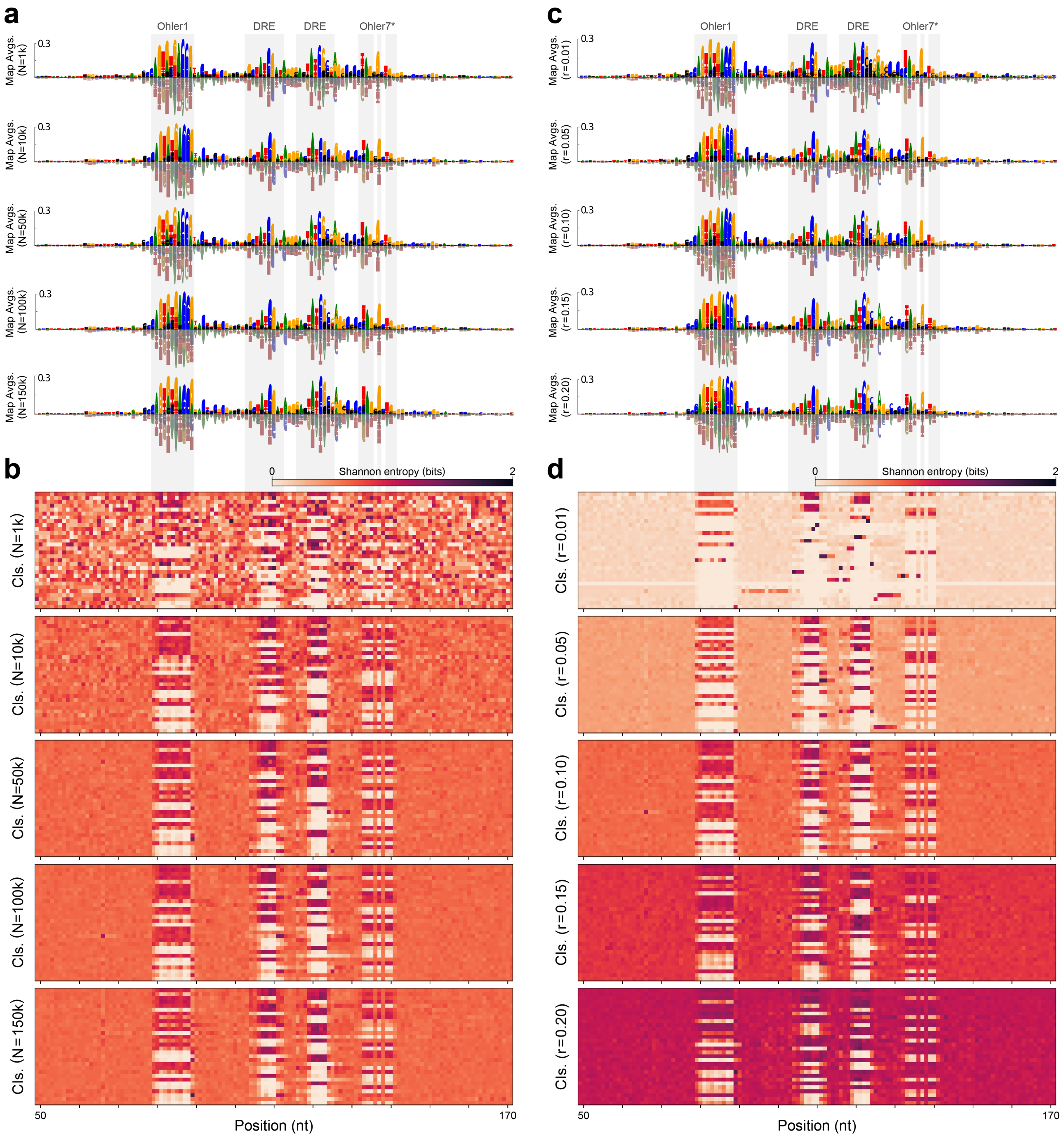


**Supplementary Figure 6. Impact of hyperparemeters on SEAM mechanistic insights.** Comparison of SEAM outputs for the DeepSTARR locus shown in Fig. 2a as SEAM hyperparameters are varied—including the size of the sequence library, $N$, and the mutation rate, $r$, used to generate the library. **a**, Comparison of SEAM variability logos—generated via the overlay of all average (avg.) attribution maps for 30 clusters (cls.) generated by hierarchical clustering—as $N$ is varied with a constant mutation rate, $r = 0.10$. **b**, Comparison of the corresponding CSMs based on positional Shannon entropy. **c**, Comparison of SEAM variability logos as $r$ is varied with a constant library size of $N = 100,000$ sequences. **d**, Comparison of the corresponding entropy-based CSMs. By visual inspection, SEAM outputs are robust to library size and rate of partial mutagenesis. Comparing the CSM shown for $N = 100,000$ and $r = 0.10$, which is a replicate of the CSM shown in Fig. 2a generated with a different random mutagenesis seed, shows that SEAM outputs are also robust to random sequence generation.

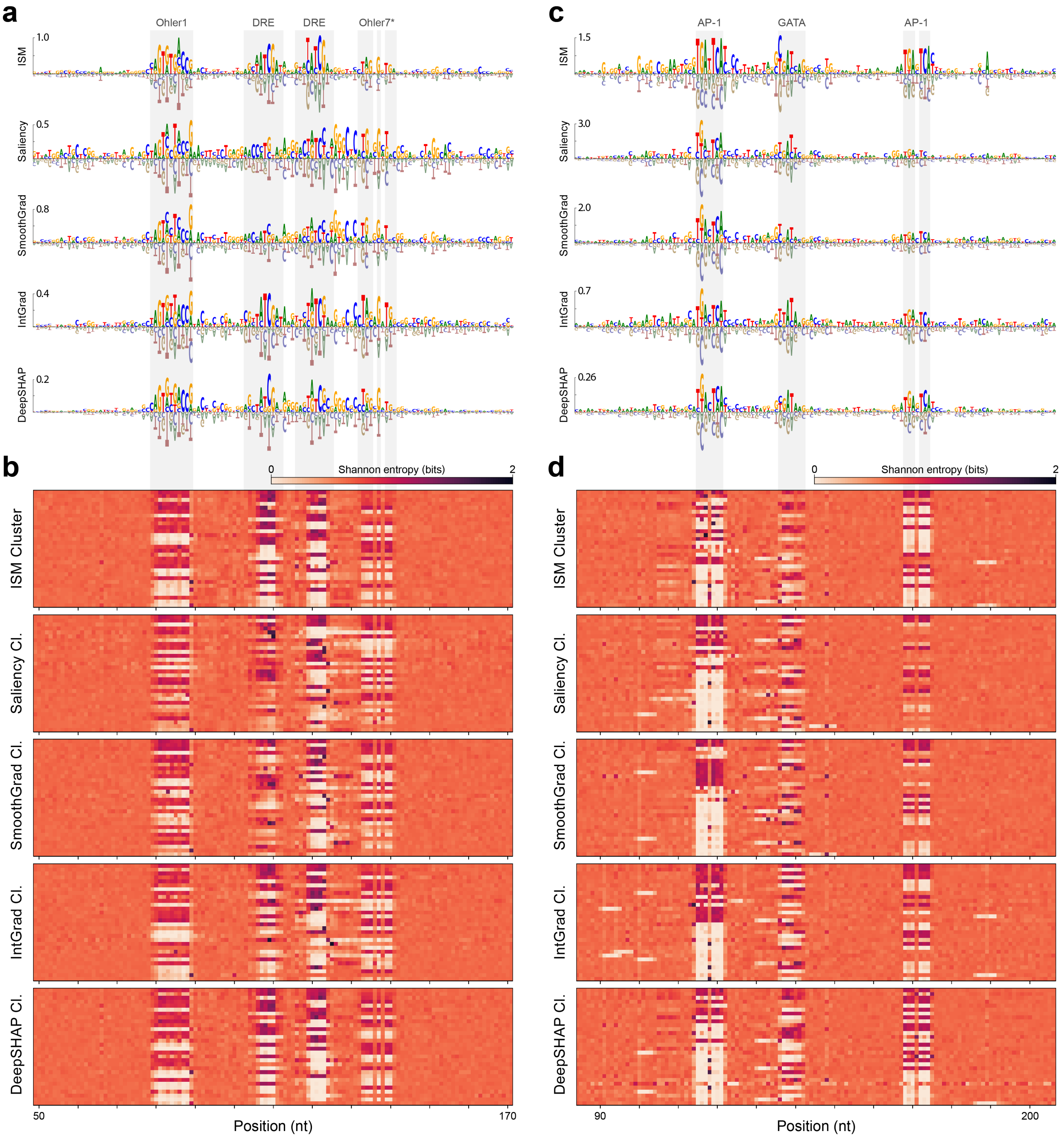


**Supplementary Figure 7. Impact of attribution method on SEAM mechanistic insights. a**, Comparison of wild-type attribution maps for the DeepSTARR locus shown in Fig. 2a, generated using *in silico* mutagenesis (ISM), Saliency Maps, SmoothGrad, Integrated Gradients (IntGrad), and DeepSHAP. Gray bars running vertically across the attribution maps align with entropy-biased regions in the corresponding CSMs, below. **b**, Entropy-based CSMs generated by SEAM using different attribution methods for the same locus. Features in the CSM are consistent across attribution methods, and identify locations of important motifs. **c,d**, Attribution maps and corresponding SEAM entropy-based CSMs for another locus obtained from the DeepSTARR test set (index 22612) using the developmental head, with similar trends observed. Cl., cluster.

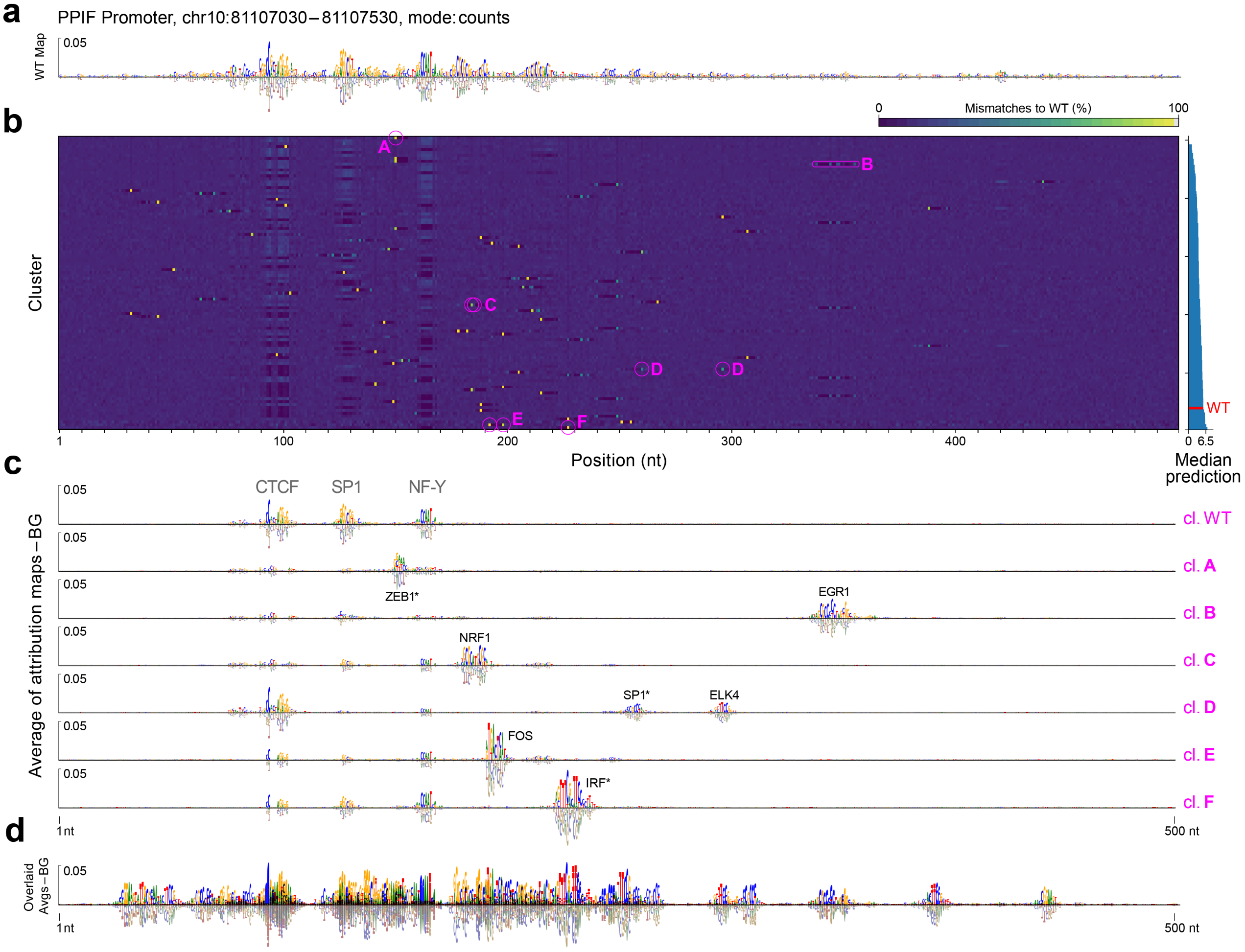


**Supplementary Figure 8. SEAM disentangles mechanistic heterogeneity at the ChromBPNet PPIF promoter. a**, Attribution map for the wild-type (WT) PPIF promoter. **b**, Cluster Summary Matrix (CSM) showing the percent mismatch at each position across sequences within a cluster, highlighting sites where mutations drive distinct mechanisms. **c**, Representative background-separated, cluster-averaged attribution maps from SEAM, each corresponding to a distinct mechanism governed by the encircled position in the CSM above (magenta). **d**, Overlay of all 100 background-separated, cluster-averaged attribution maps reveals the full landscape of mechanistic diversity across the local library.

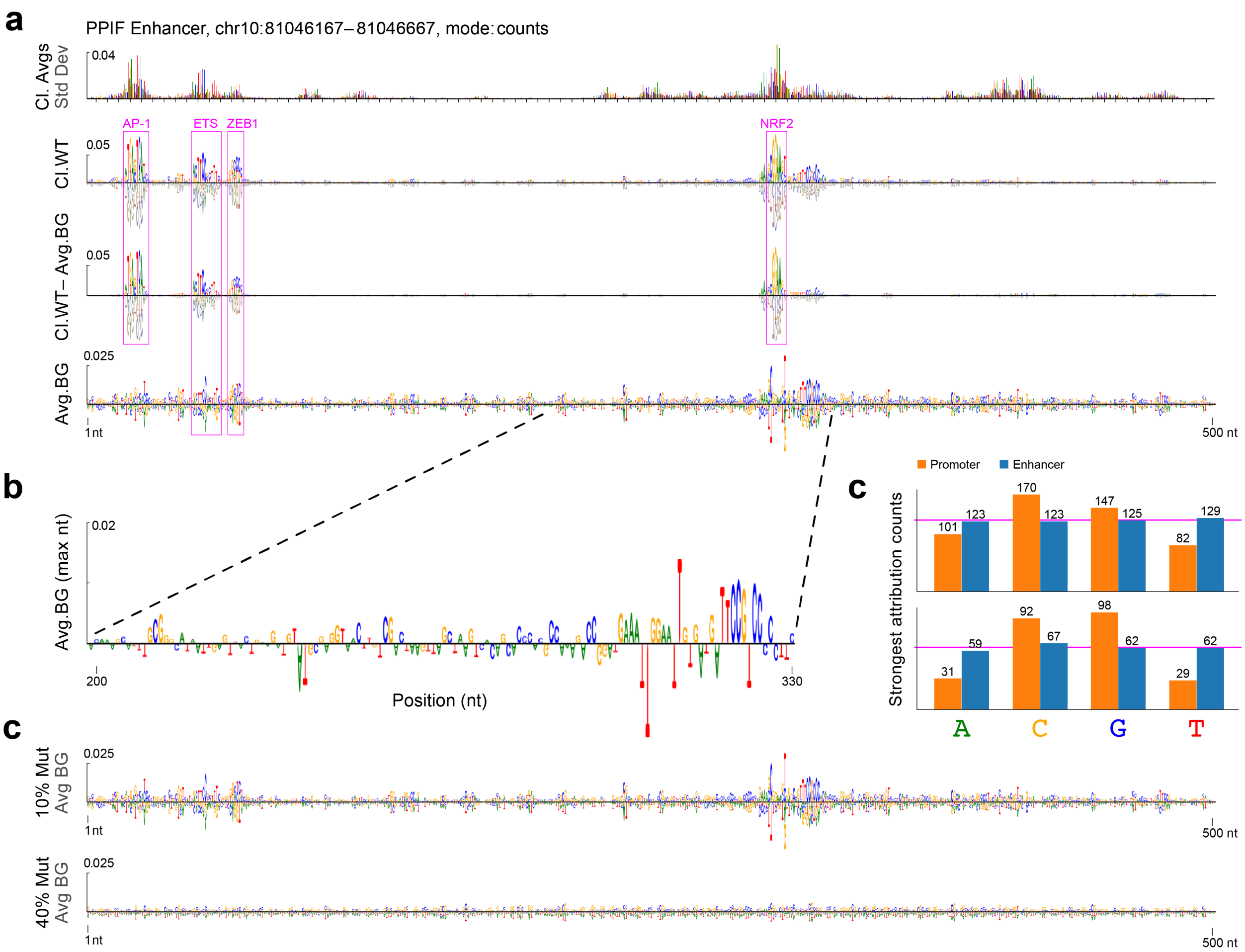


**Supplementary Figure 9. SEAM-defined background signals reveal compositional and mutational differences between the PPIF promoter and enhancer. a**, Standard deviation of attribution values across all clusters at the PPIF enhancer (10% mutation rate using the first model fold), illustrating the static background on which TF motifs reside. **b**, Attribution analysis of the enhancer at 10% mutation. Top: attribution map for the wild-type cluster (cl.WT). Bottom: SEAM-derived average background (Avg. BG) computed across clusters. Middle: Result of subtracting the average background from the WT cluster. While AP-1 and NRF2 motifs are ablated in the background and appear only in the WT cluster, residual attribution signals corresponding to ETS and ZEB1 motifs persist following SEAM's background separation, indicating partial resistance to 10% mutational perturbation. **c**, Visualization of the enhancer's average background attribution map (from **b**, bottom) displaying only the most prominent nucleotide at each position (maximum attribution), highlighting its compositionally mixed signal. **d**, Comparison of nucleotide attribution patterns between the PPIF promoter (orange) and enhancer (blue). For each position, the nucleotide with the strongest absolute attribution was identified. Top: Distribution across all positions. Bottom: Restricted to the top 50% of positions by attribution magnitude. In both cases, promoter attributions are skewed toward C/G, while the enhancer exhibits balanced nucleotide composition. Pink horizontal lines denote perfectly uniform distribution. **e**, SEAM-derived average background attribution maps at the PPIF enhancer under 10% (top) and 40% (bottom) mutation rates. At 40%, all motif signals are lost, leaving only low-attribution, diffuse background signals.

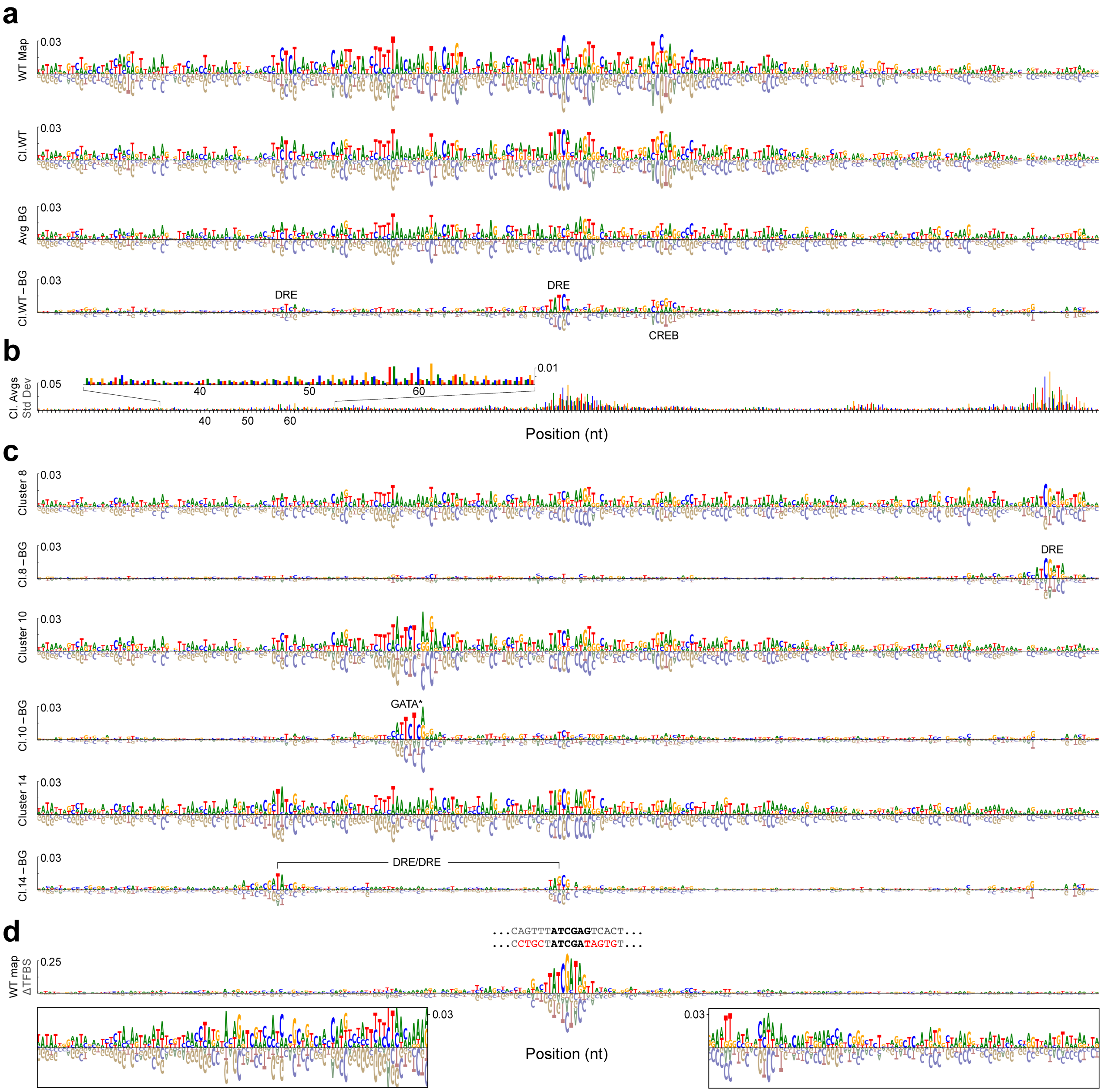


**Supplementary Figure 10. Background separation at a DeepSTARR enhancer.** **a**, First row: Wild-type (WT) attribution map from a DeepSTARR enhancer (test set index 4071). A low-affinity DRE TFBS, ATCGAG–with one mutation to the consensus TFBS, ATCGAT–is positioned near the center, resulting in an overall low Hk expression (-0.49). Seemingly spurious scores across the attribution map obscure the identification of TF motifs. Second row: Average (avg.) attribution map for the WT cluster (cl.WT), showing SEAM's ability to denoise the WT mechanism by averaging over qualitatively similar attribution maps. Third row: Average of all intra-cluster backgrounds (BG) separated by SEAM, featuring AT-rich attribution signals across the enhancer. Fourth row: WT cluster after background removal, generated by subtracting the average background (row 3) from the averaged WT mechanism (row 2). Background separation reveals three previously obscured TF motifs. **b**, Standard deviation of attribution values over clusters, highlighting the static nature of the background on which the TF motifs reside across all clusters. **c**, Examples of background separation on other mechanisms discovered by SEAM at the same locus, revealing previously obfuscated TF motifs in each cluster. Cluster 14 recovers a higher-affinity DRE/DRE motif compared to WT that is not necessitated by CREB on the right-hand side. **d**, Attribution map was generated from the WT sequence after mutating the central DRE TFBS to match the consensus motif ATCGAT, including optimized flanking nucleotides. Compared to the WT, these 10 mutations increase Hk expression to 2.01, while substantially altering the attributed background context across the enhancer (see insets). This example highlights the sensitivity of background signals to coordinated mutations.

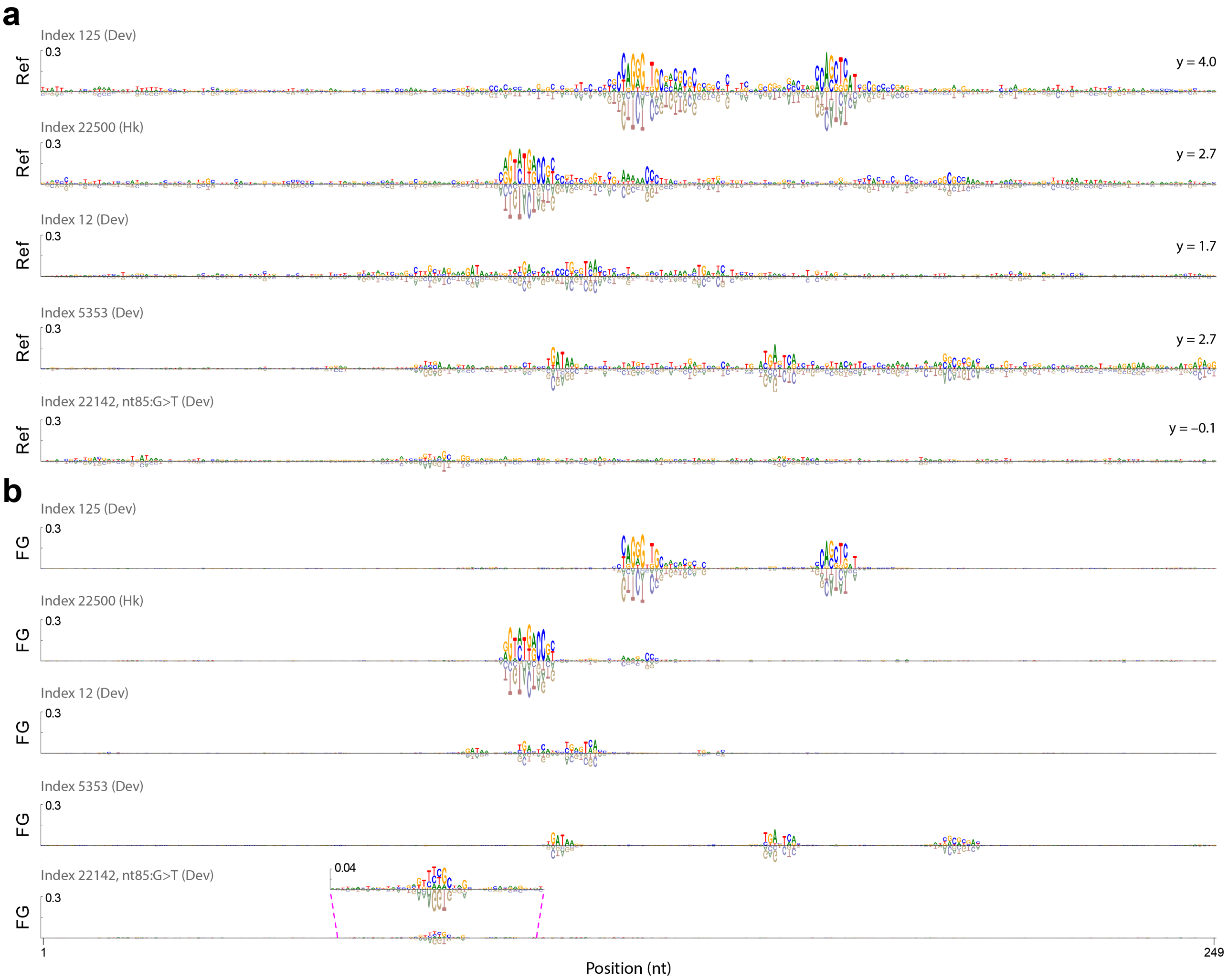


**Supplementary Figure 11. SEAM background separation enhances motif clarity across DeepSTARR enhancers.** **a**, Each panel shows the DeepSHAP attribution map for a reference (Ref) sequence curated from the DeepSTARR test set. DeepSTARR predictions ($y$) for each sequence are listed on the right. **b**, SEAM foreground (FG) attribution map computed for each reference sequence (see Methods). Background separation was derived using SEAM from a locally mutagenized library (10% mutation rate, 100k sequences, 30 clusters). Following SEAM-based clustering and background separation, reference motifs emerge with striking clarity, revealing the underlying regulatory logic that was previously obscured. Dev, developmental head; Hk, Housekeeping head.

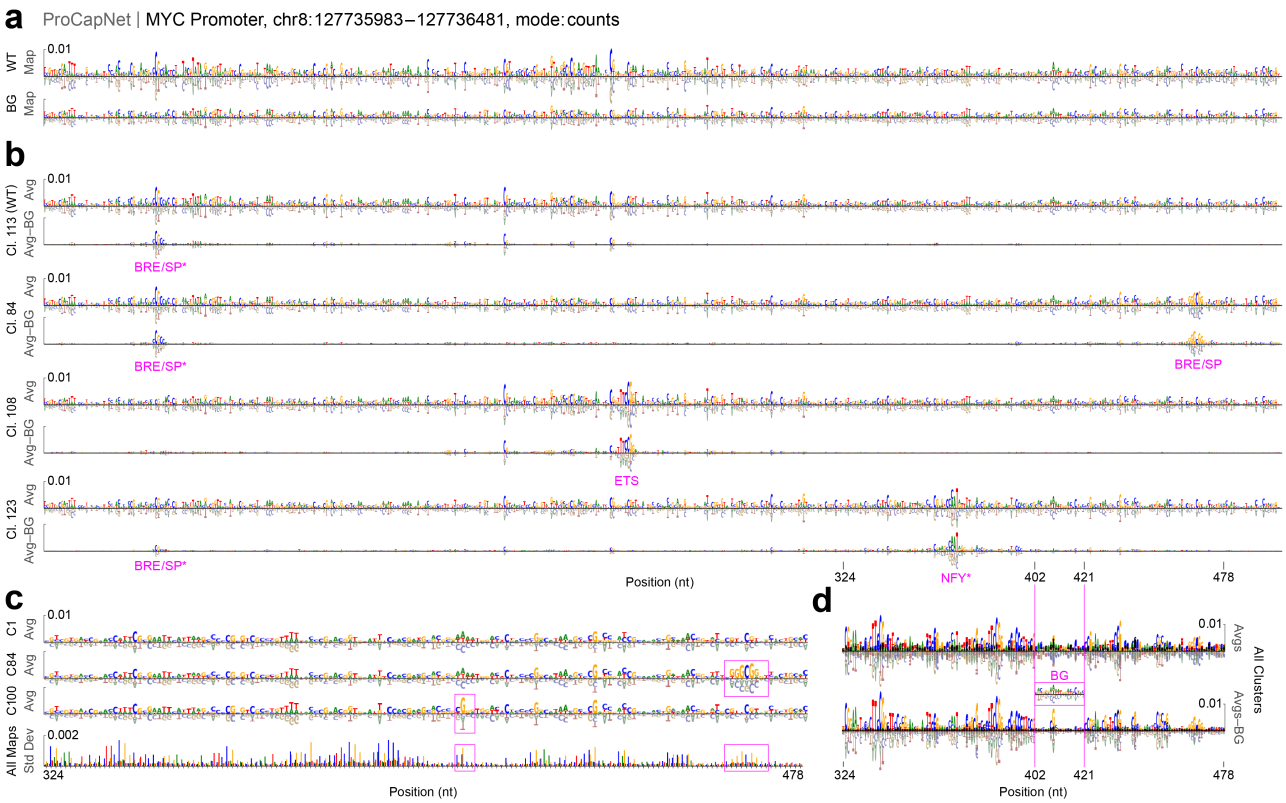


**Supplementary Figure 12.** **Background separation at the ProCapNet MYC promoter.** **a**, Top: Wild-type (WT) attribution map with seemingly spurious scores across the promoter, obscuring the identification of TF motifs. Bottom: Average (avg.) background (BG) across all clusters, as separated by SEAM. **b**, Examples of four clusters out of 200, each showing before (top) and after (bottom) background separation on the average attribution maps in the associated cluster. **c**, Examples of averaged attribution maps from three clusters, showing the persistence of background signals across mechanisms. While the standard deviation over clusters is relatively large at the positions of mechanism-specific motifs (highlighted in pink rectangles), the background attribution signals surrounding them show nearly zero standard deviation, emphasizing the static nature of the background on which the TF motifs reside across all clusters. **d**, Overlay of the average attribution maps in all 200 clusters before (top) and after (bottom) background separation. Cl./C., cluster.

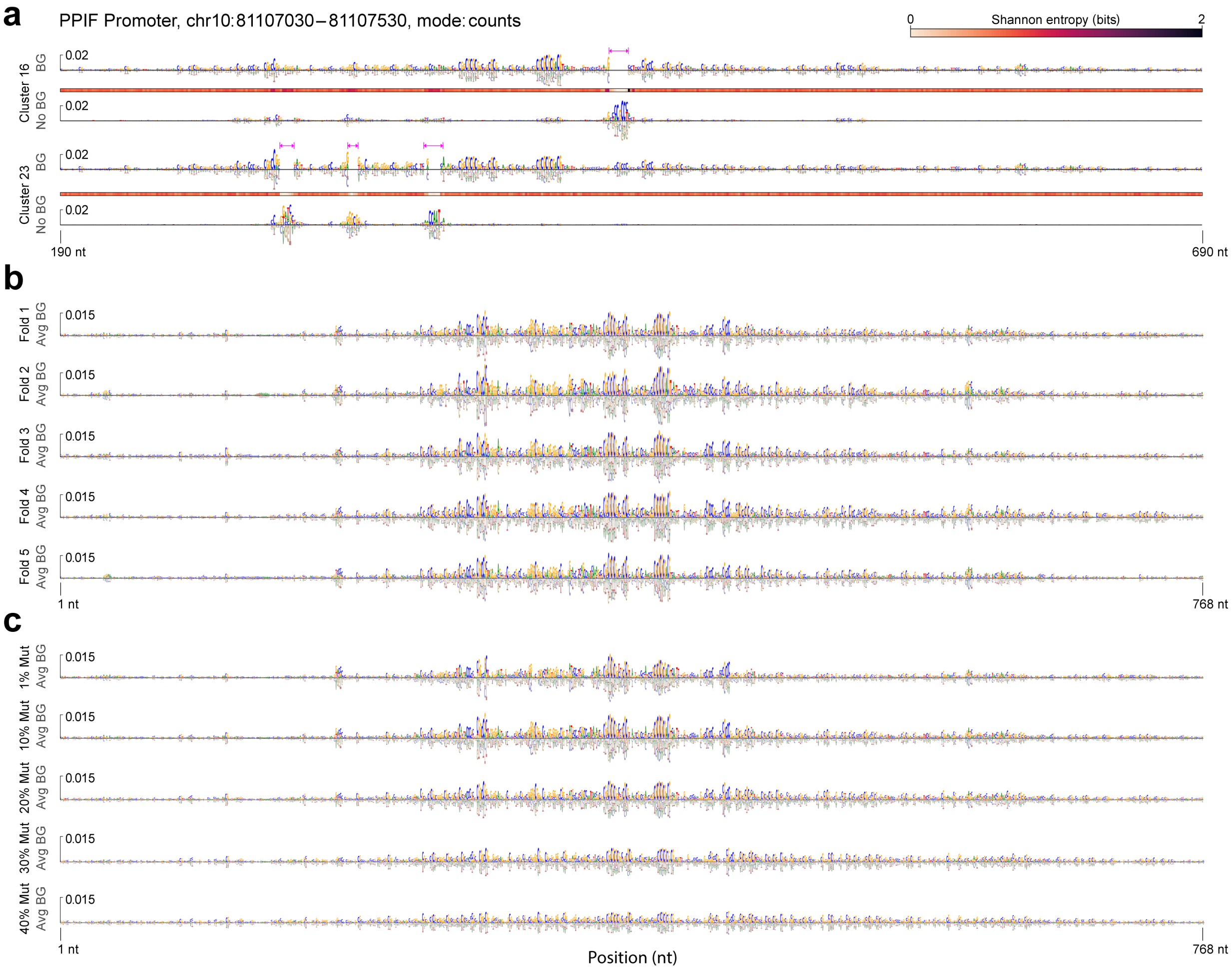


**Supplementary Figure 13. Impact of ChromBPNet model fold and mutation rate on SEAM background separation.** **a**, Examples of attribution maps for intra-cluster background (BG, top) and foreground (FG, bottom) for clusters 16 and 23. FG is defined by subtracting the average of all intra-cluster BGs from the average of attribution maps in a given cluster. Intra-cluster background is informed by the positional Shannon entropy over sequences in each cluster (middle). SEAM uses the thresholded positional Shannon entropy to mask out TF-dependent attributions in each map. The foreground attribution map is derived by subtracting the average (avg.) attribution map of a given cluster from the average of all intra-cluster BG maps over all clusters. As in Fig. 3, SEAM was run using the average of attribution maps over all folds, with sequences sampled using a 10% mutation rate. **b**, Results of running SEAM background separation independently on each of the first five ChromBPNet folds (10% mutation rate). **c**, Results of independently running SEAM background separation on the first fold using datasets generated with different mutation rates.

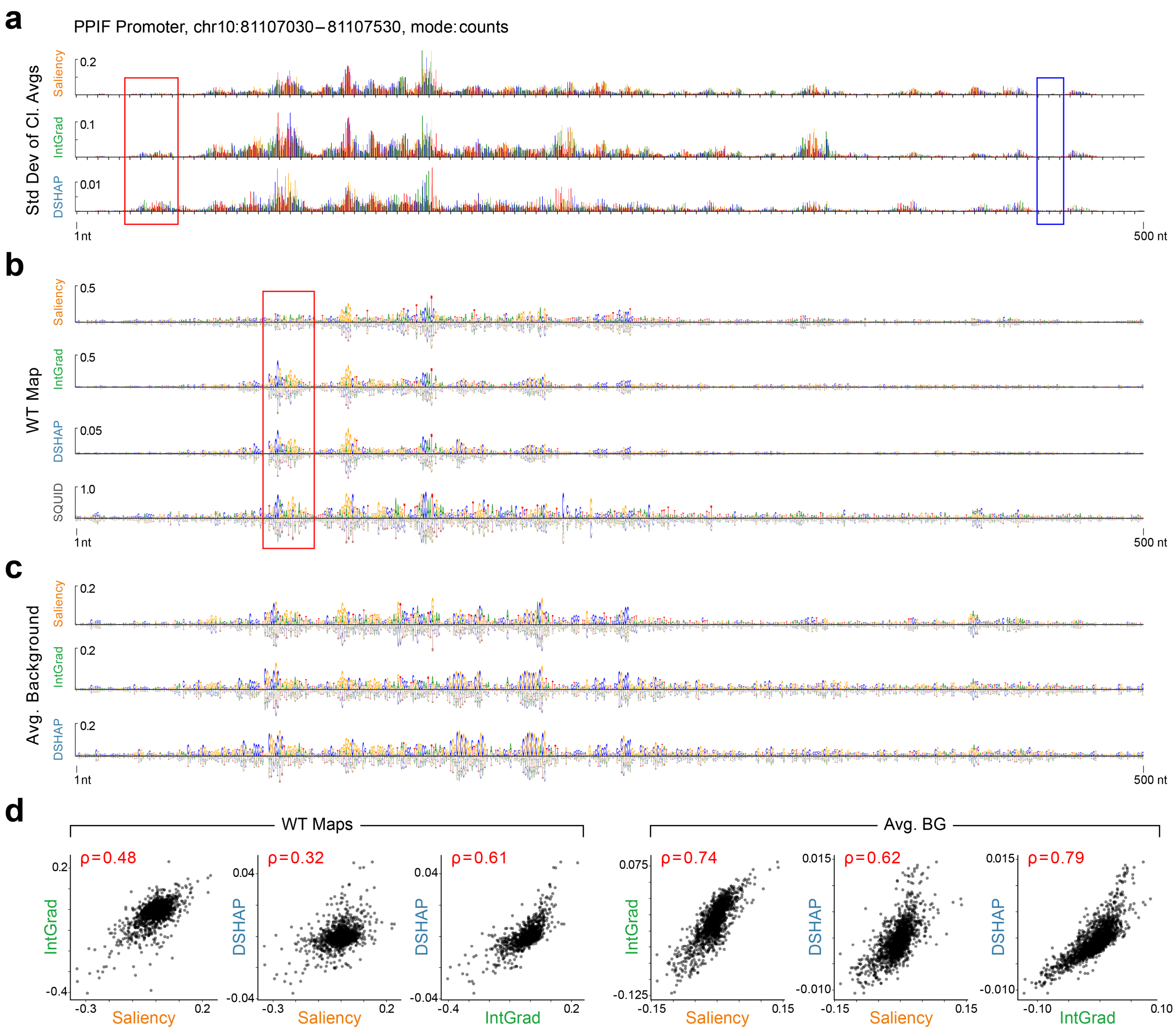


**Supplementary Figure 14. SEAM reveals consistent background signals across attribution methods despite divergent foreground quality. a**, Standard deviation of cluster-averaged attribution maps at the ChromBPNet PPIF promoter for Saliency Maps (Saliency), Integrated Gradients (IntGrad), and DeepSHAP (DSHAP). For each attribution method, SEAM was run independently on the same library of 100,000 sequences generated with a 10% mutation rate, using the first model fold, and clustered using hierarchical clustering into 200 clusters. Saliency exhibits the highest overall variation across positions, followed by IntGrad and then DeepSHAP. Standard deviation profiles are broadly consistent across methods, with shared regions of near-zero variability (e.g., blue box) suggesting convergence on background-dominated signals. However, a motif-length region shows a notable discrepancy (red box), where Saliency displays minimal variability while IntGrad and DeepSHAP detect a poised motif. **b**, Attribution maps for the WT PPIF promoter computed using Saliency, IntGrad, DeepSHAP, and SQUID. SQUID was trained on the same local neighborhood as SEAM and achieved an additive model fit of $R^2 = 0.54$. A key motif absent in Saliency but recovered by other methods is highlighted (red box). Relative to IntGrad and DeepSHAP, Saliency lacks a clearly defined GC-rich background, suggesting reduced sensitivity to local regulatory context. **c**, Average background maps separated by SEAM for Saliency, IntGrad, and DeepSHAP. Despite differences in foreground quality, SEAM isolates highly similar background signals across methods, consistent in both spatial pattern and attribution magnitude. **d**, Left: Pairwise scatter plots comparing WT attribution maps across methods (Saliency vs IntGrad, Saliency vs DeepSHAP, IntGrad vs DeepSHAP), with Spearman correlation coefficients shown. Right: Corresponding scatter plots comparing SEAM-separated average background maps across methods. Correlation values increased by 0.26, 0.30, and 0.18, respectively (from 0.48, 0.32, and 0.61 to 0.74, 0.62, and 0.79), indicating that background signals are substantially more consistent across attribution methods than the raw attribution maps themselves.

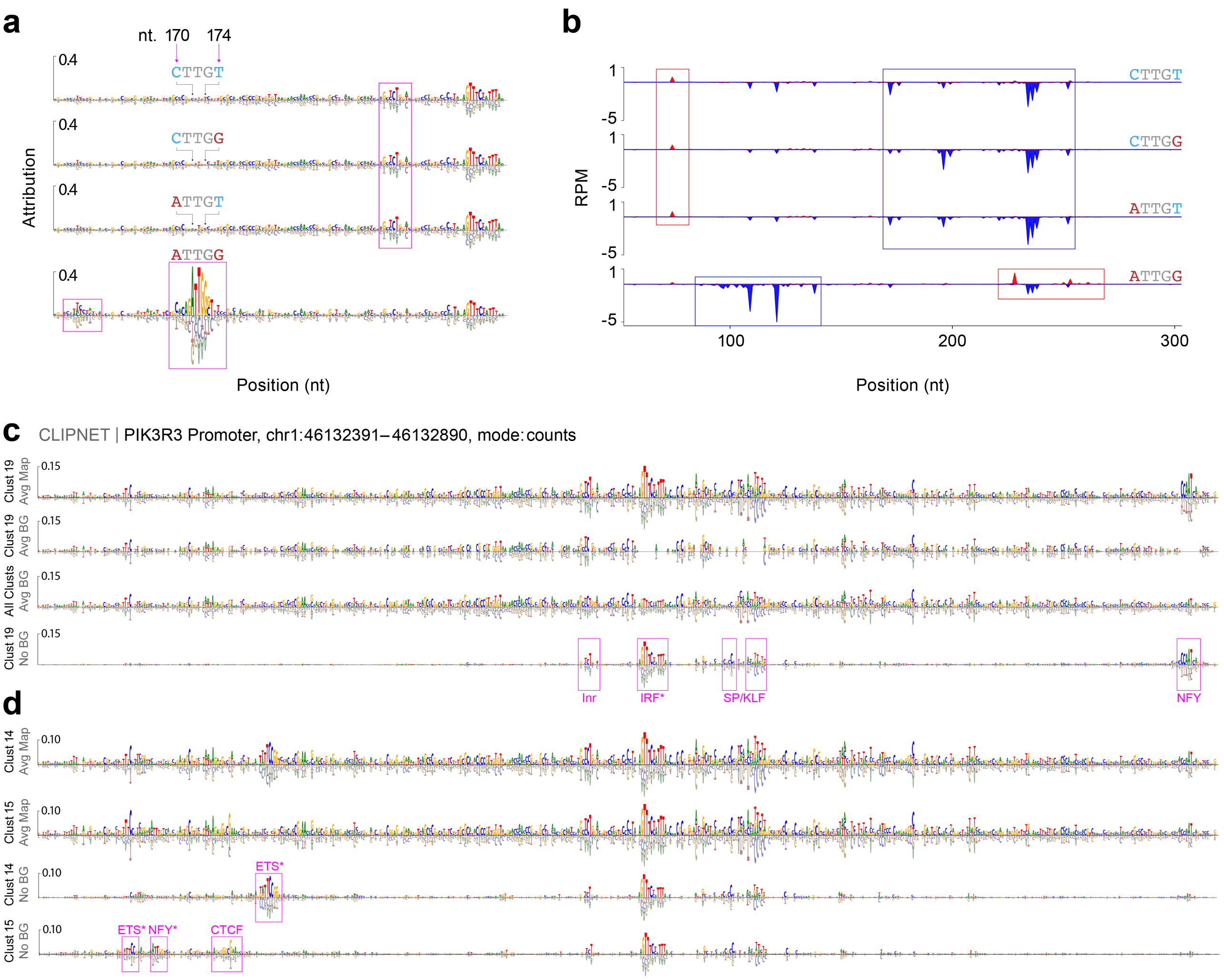


**Supplementary Figure 15. Attribution analysis at the CLIPNET PIK3R3 promoter. a**, CAAT box TF motif analysis using attribution maps generated from sequences with none, one, or both of the SEAM-discovered pairwise mutations (cl. G in Fig. 4). In the last row, both mutations are applied to the WT sequence, revealing a CAAT box binding motif along with upstream and downstream mechanistic changes. **b**, CLIPNET predicted profile for the four sequences in the CAAT box analysis, showing that SEAM-discovered mutations change transcription direction at the promoter. As a general note on the computational challenge of capturing pairwise effects, the number of double nucleotide variants ($k = 2$) at the CLIPNET promoter (length $L = 500$, $A = 10$ alleles) is $\binom{L}{k} \cdot (A-1)^k$, requiring assessment of 10,112,251 attribution maps. RPM, reads per million. **c**, Mechanism-decomposed attribution maps previously discovered at the CLIPNET PIK3R3 promoter (in Fig. 4) can be further decomposed to separate background from TF-dependent motifs. First row: The average (avg.) attribution map for cluster 19 (cl.19). Second row: The intra-cluster average background for cluster 19, with masking informed by the positional Shannon entropy over sequences in the cluster. Third row: The average of all intra-cluster backgrounds (BG) from each cluster. Fourth row: Subtracting the third row from the first row produces a noise-reduced, background-separated representation of the mechanism in cluster 19. **d**, Additional examples of SEAM noise-reduced background separation for clusters 14 and 15.

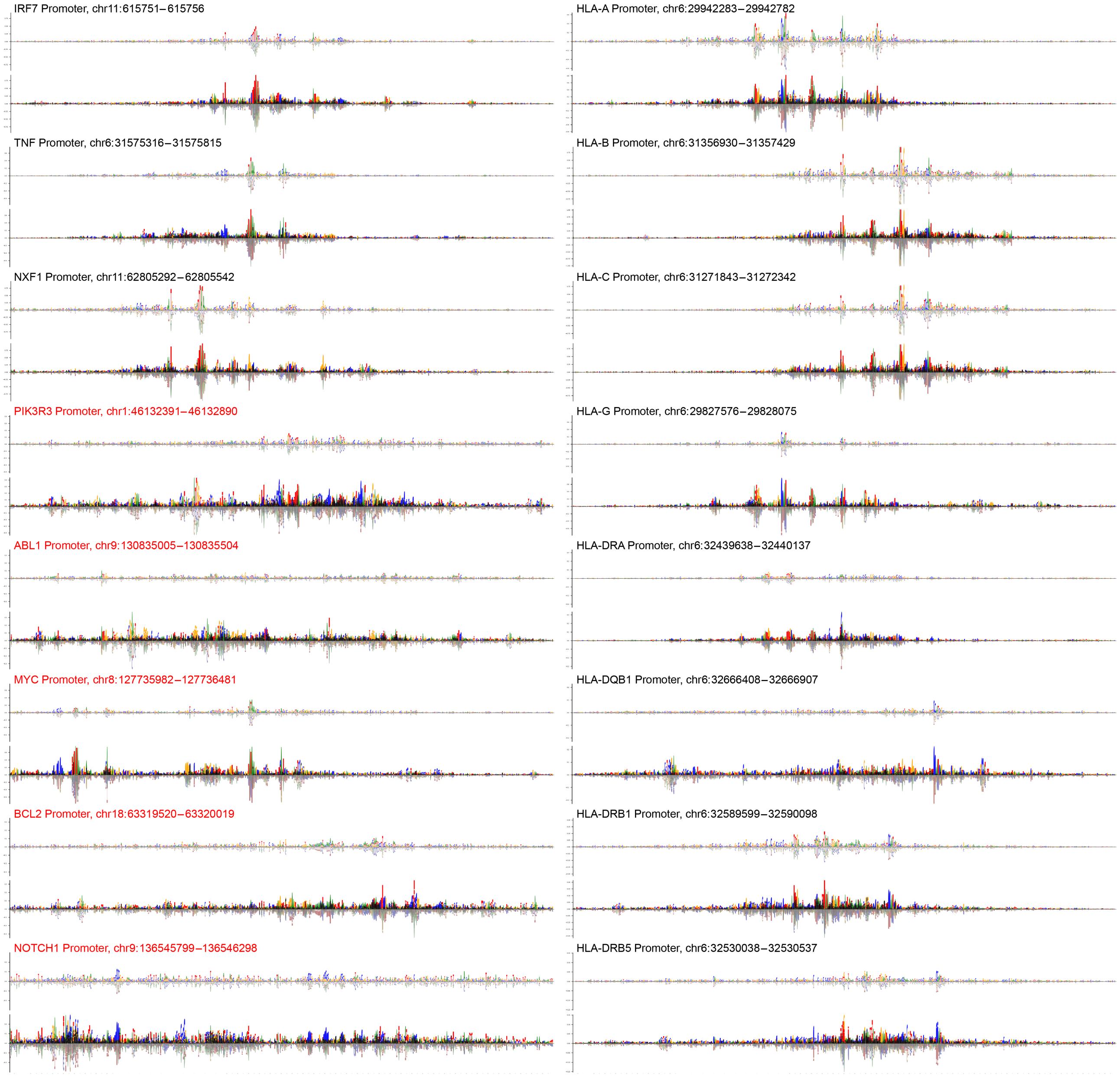


**Supplementary Figure 16. SEAM analysis of mechanistic variation at CLIPNET promoters.** Wild-type (WT) attribution map generated by CLIPNET (top) and the overlay of all average attribution maps for the 200 clusters (bottom), generated by SEAM, for each of sixteen promoters. Non-oncogene and oncogene promoters are denoted with black and red text, respectively. A subset of the curated promoters are highly polymorphic, which includes the oncopromoters and HLA family (right).[1] In non-oncogene promoters, mechanistic variation is typically constrained to tuning the amplitude of constitutively active motifs already present in the WT attribution map. In these cases, the mechanistic variation discovered by SEAM is highly conserved. In contrast, oncogene promoters show substantially more mechanistic variation compared to their WT attribution map, with many poised regulatory motifs spanning the promoter sequence. SEAM also identifies conservation of mechanistic variation across related loci, such as HLA-B and HLA-C, which have a 89.2% sequence similarity as calculated by EMBOSS.[2] Beyond sequence similarity, SEAM is able to consider the similarity between mechanistic aspects of promoter regulation.

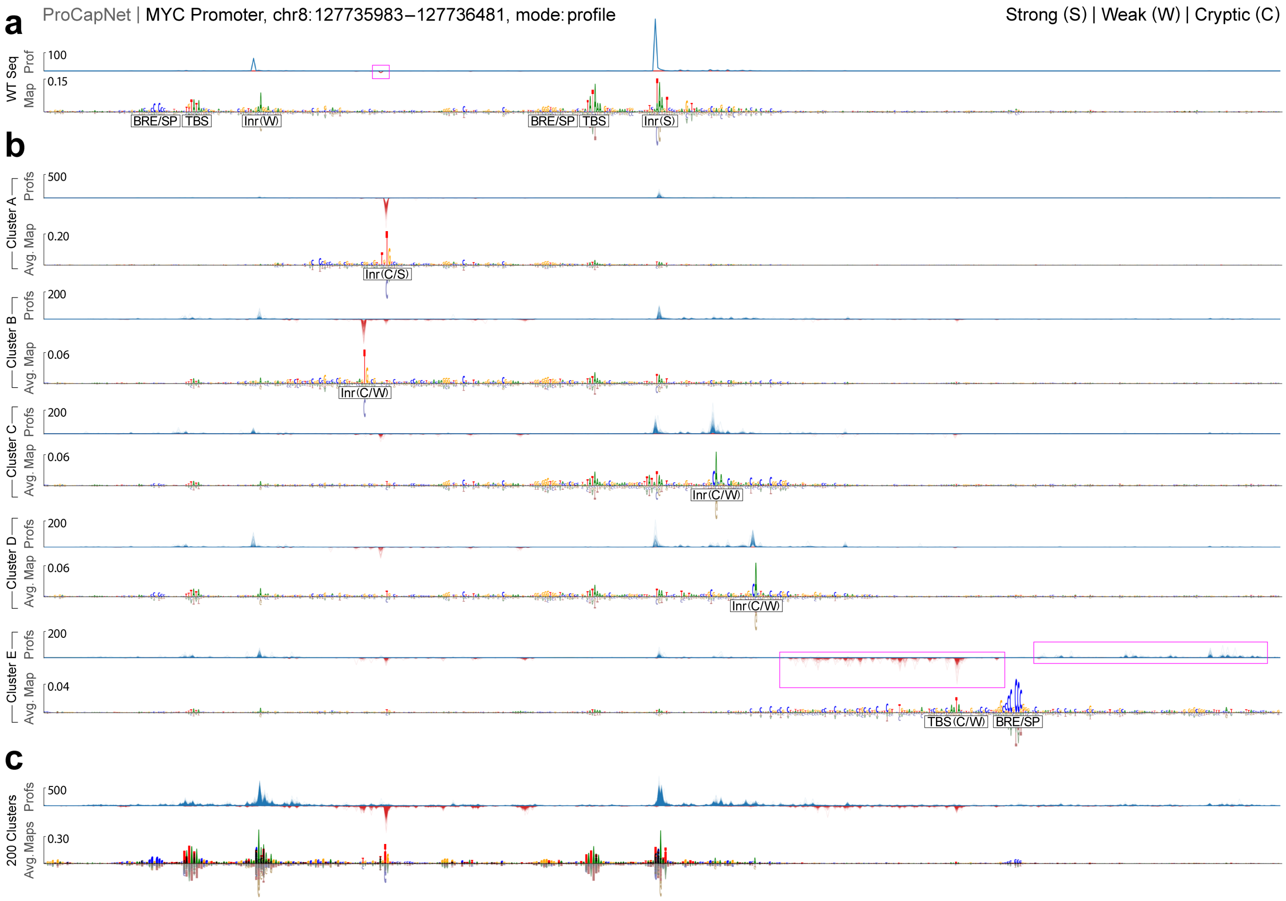


**Supplementary Figure 17. SEAM analysis of mechanistic variation at ProCapNet MYC promoter.** **a**, Predicted profile and attribution map for the wild-type sequence (WT Seq), demonstrating the TF motifs and profile peaks present, which are used for biased ablation experiments. **b**, Five (cl. A-E) of the 200 clusters generated using SEAM. For each panel, the predicted profiles (prof.) associated with the attribution maps in the given cluster are overlaid on top, with the average (avg.) of all attribution maps in the cluster displayed on the bottom. In cluster A, SEAM finds a stronger version of the previously-documented weak antisense initiator. In clusters B-D, SEAM finds previously-undiscovered weak antisense (cl. B) and sense (cl. C, D) initiators. In cluster E, SEAM discovers an alternative TATA and BRE/SP TFBS that substantially alters upstream and downstream attribution values, while reversing the direction of transcription (pink rectangles). **c**, Overlay of all predicted profiles for all clusters (top) and overlay of all average attribution maps per cluster (bottom). In this view, the appearance of smeared motifs represent overlapping mechanisms that can be finely tuned over a broad range based on sequence context.

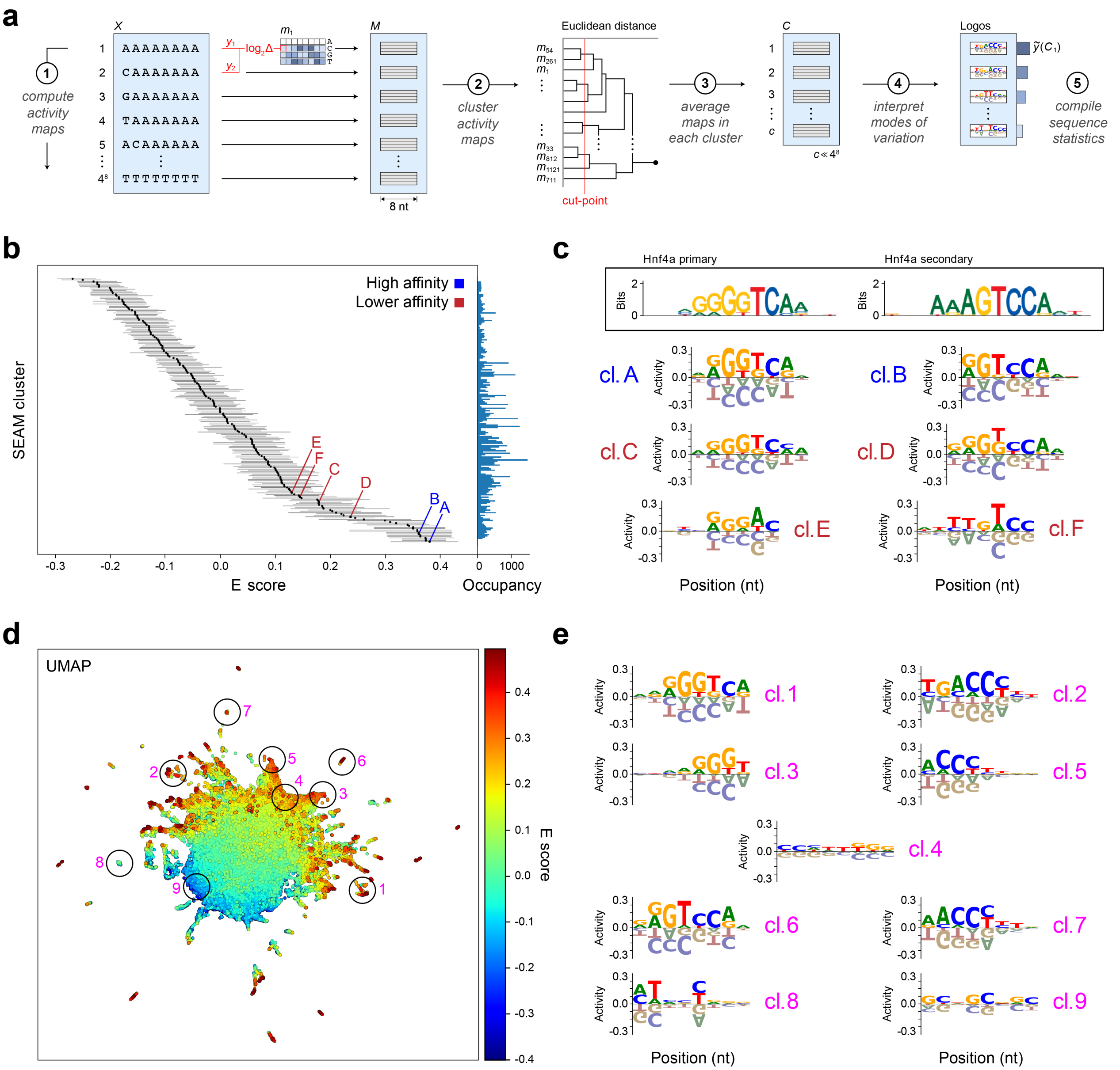


**Supplementary Figure 18.** **SEAM identifies alternative binding preferences of Hnf4a.** **a**, Schematic for applying the SEAM framework to combinatorially complete libraries. For every sequence in the experimentally obtained Protein Binding Microarray (PBM) library, SEAM generates an empirical mutagenesis map using the log2 fold change between the E scores of the reference sequence and a single nucleotide variant. All other steps in the SEAM framework remain unchanged. **b**, Left: Box plots showing E scores for each of the 200 clusters produced by SEAM, using hierarchical clustering, ranked in ascending order by each cluster's median E score. Lines represent the upper and lower quartiles; the average IQR across clusters is 0.10. Letters A-F label example clusters. Right: Number of empirical mutagenesis maps (occupancy) in each cluster. **c**, Top inset: Information content sequence logos for the two alternative binding modes captured in the original PBM study of Hnf4a using the Seed-and-Wobble algorithm. Bottom: Sequence logos of averaged empirical mutagenesis maps corresponding to the example SEAM clusters labeled in the previous panel. **d**, UMAP embedding of the 100,000 attribution maps, as used in previous panels, where each attribution map is represented as a point colored by the E score of its corresponding sequence. Numbers 1-9 label example clusters. **e**, Sequence logos for empirical mutagenesis maps averaged within each encircled regions in the previous panel.

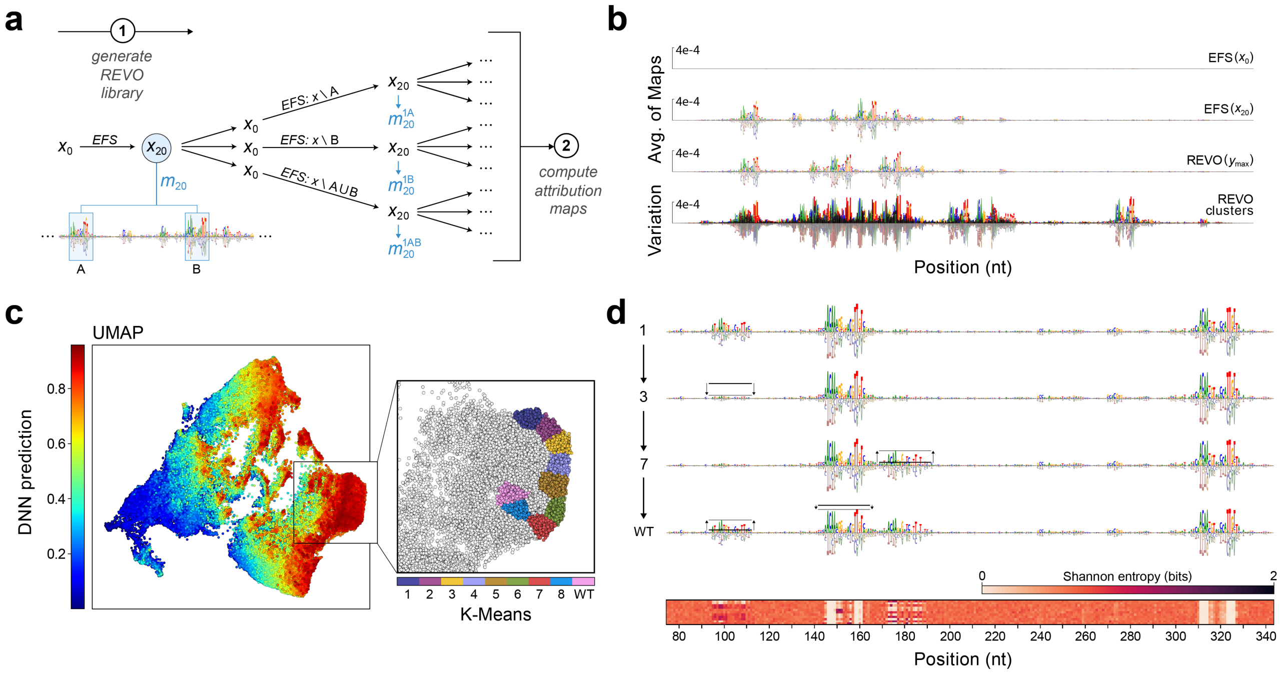


**Supplementary Figure 19.** **SEAM mechanistically fine-tunes pre-optimized sequences.** **a**, Schematic of the REVO protocol for designing mechanistically diverse sequences optimized for a given objective (see Methods). **b**, First row: Attribution map for the EFS-6 starting sequence, obtained via the Evolved from Scratch algorithm. Second row: Attribution map for the final sequence in the EFS-6 trajectory. Third row: Attribution map for the highest-scoring REVO sequence initialized from EFS-6, which is qualitatively distinct from the second row. Fourth row: Overlay of average maps from each of the 200 clusters generated by SEAM, applied to the REVO library initialized from EFS-6. **c**, UMAP embedding of attribution maps, where each attribution map is represented as a point colored by DNN prediction. Attribution maps were generated from a local library (100,000 sequences using a 10% mutation rate) using the final sequence from the EFS-4 pathway. The resulting mechanism space uncovers a continuum of attribution maps organized by DNN prediction. After clustering these maps into 200 groups using K-means, we identified distinct trajectories across contiguous clusters that fine-tuned the contributions of individual motifs in relation to their roles in the initially optimized mechanism. An example trajectory formed from nine clusters is shown in the inset. **d**, Top: Averaged attribution maps of the clusters shown in the previous inset. Following the trajectory across clusters fine-tunes the final EFS-4 mechanism. Bottom: Subset of the CSM based on positional Shannon entropy, highlighting the unique sequence determinants driving mechanistic variation in mechanism space. Analysis of the CSM reveals sequence dependencies responsible for fine-tuning these motifs are idiosyncratic and unpredictable from sequence alone.

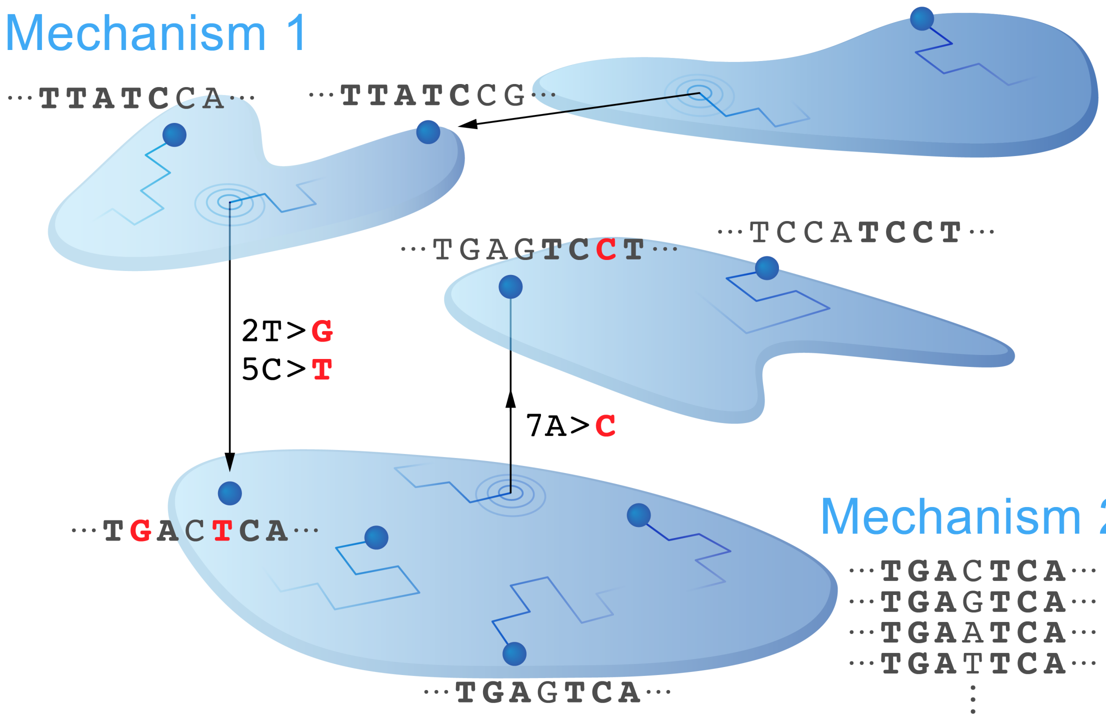


**Supplementary Figure 20. Schematic of the terraced labyrinth fitness function.** Schematic of Crutchfield and Nimwegen's terraced labyrinth fitness function[3], illustrating mutational trajectories constrained within mechanisms (subbasins) and rare portal mutations that drive shifts between them. This subbasin–portal architecture captures epochal evolution, with long periods of stasis punctuated by sudden innovations.

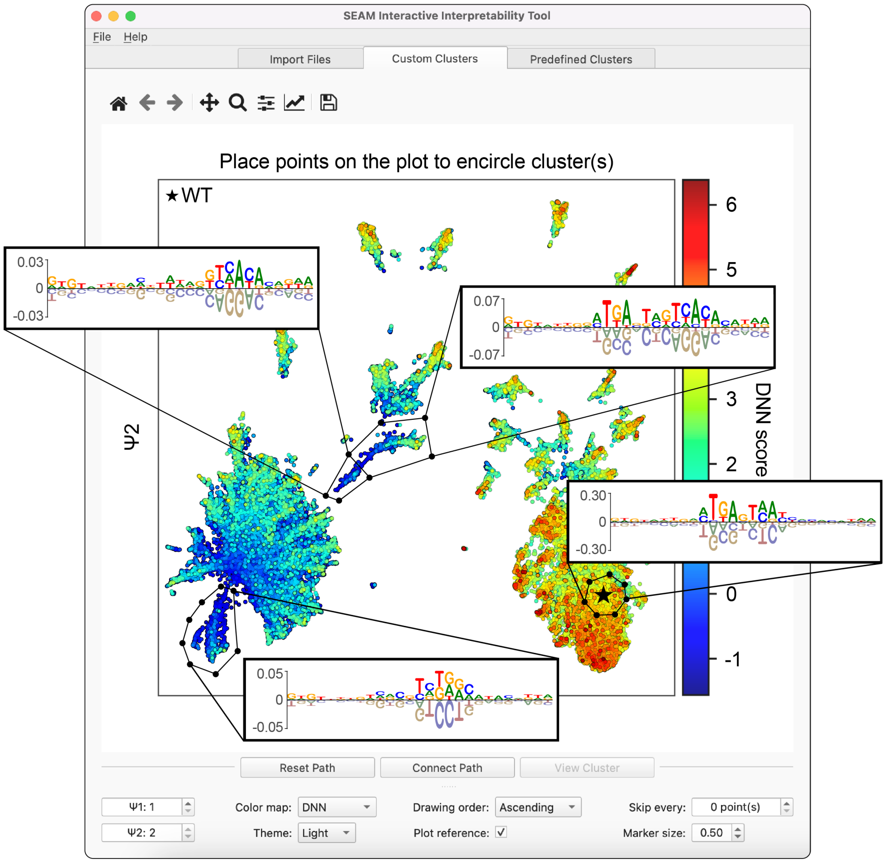


**Supplementary Figure 21. SEAM Graphic User Interface (GUI).** Example usage of the SEAM Interactive Interpretability Tool, a GUI allowing users to explore SEAM outputs, such as defining clusters with customized boundaries in the embedded space (shown) and analyzing associated sequence statistics. In the embedded space, each point is an attribution map from a local library, colored by its predicted activity. Attribution maps and related statistics in user-drawn clusters are averaged and displayed in a variety of formats. Data in schematic generated using DeepSTARR: test set index 13748; developmental head.

### Supplementary Note 1: Relation of SEAM to other methods

SEAM was developed to characterize the diversity of regulatory mechanisms accessible through local mutational variation and separate foreground from background regulatory signals. While related in spirit to several interpretability approaches, SEAM differs in scope and strategy. Below, we highlight conceptual distinctions from prior work.

**Motif discovery frameworks**

- **TF-MoDISco**[4] clusters high-attribution segments from large-scale genomic data to extract recurring sequence motifs. These motif-level summaries are effective for cataloging learned features but are not designed to resolve multiple mechanistic programs at a single locus. SEAM instead focuses on local neighborhoods of sequence space, constructing mutagenized libraries aligned to a reference and clustering full-length attribution maps. This enables the discovery of distinct regulatory mechanisms, the specific mutations that induce them, and the separation of foreground from background regulatory signals—capabilities not addressed by motif discovery tools.

**Attribution smoothing approaches**

- **SmoothGrad**[5] improves attribution interpretability by averaging maps across inputs perturbed with Gaussian noise. While effective in vision models, this strategy moves genomic sequences off the discrete DNA alphabet, undermining biological validity. SEAM generalizes this denoising strategy using biologically meaningful perturbations—point mutations—and ensures averaging occurs only among sequences that share a consistent mechanistic program. As such, SEAM simultaneously reduces mechanism-specific attribution noise while revealing the distinct mechanisms poised near a given sequence.

**Model-guided perturbation methods**

- **Counterfactual motif ablation** experiments focus narrowly on mutating putative motifs and recalculating attribution maps to assess changes in regulatory mechanisms.[6] While useful for validating the importance of known features, these approaches overlook the contribution of nucleotides outside of easily visualized motif patterns or regions with elevated signal in predicted profile tracks. As a result, they provide a limited and potentially biased view of mechanistic diversity, missing context-dependent or distributed regulatory features. In contrast, SEAM samples a broader, unbiased mutational neighborhood, enabling the discovery of unexpected, noncanonical, or context-specific mechanisms.
- ***In silico* optimization** approaches formulate the problem of regulatory design as an objective-driven search through sequence space. For example, Ledidi performs gradient-based optimization to design discrete edits that induce a desired change in model output.[7] While well-suited for engineering specific functional outcomes, such methods assume a fixed design objective and do not explore alternative regulatory architectures. SEAM differs in scope: when using local libraries, it provides an unbiased, non–goal-directed view of mutational space; when applied to optimization libraries (e.g., REVO), SEAM clusters the resulting sequences to expose mechanistic heterogeneity within a defined region of sequence space, revealing distinct regulatory strategies that satisfy the same functional objective.

**Limitations of interpreting attribution scores as regulatory mechanisms**

- ***In Silico* Mutagenesis (ISM)** quantifies the effects of individual mutations by computing prediction differences.[8] However, high-amplitude ISM scores may reflect mutational sensitivity without indicating how the mutation rewires regulatory logic. Moreover, mechanistic interpretation requires manual inspection and targeted mutation, introducing bias toward large-effect variants while missing subtle mechanism-altering changes masked by background signal (e.g., Fig. 1c).
- **Complete and linear attribution methods** such as Integrated Gradients[9] and DeepSHAP[10] distribute importance across input features under completeness constraints. Recent theoretical work has shown that these attribution methods and others relying on linearity and completeness constraints can assign zero attribution to positions that nonetheless impact model predictions.[11]

SEAM avoids these limitations by offering an unbiased, data-driven framework for discovering mutationally induced shifts in regulatory mechanism without relying on prior assumptions about which positions or variants to test. Because attribution methods operate on a fixed input, they are inherently limited to interpreting individual sequences—failing to capture combinatorial effects of mutations or the broader structure of mechanistic variation across sequence space. By analyzing a localized mutational neighborhood and clustering full-length attribution maps, SEAM resolves distinct mechanisms, identifies their sequence determinants, and reveals how regulatory logic can be reprogrammed through mutation.

### Supplementary Note 2: Global sequence libraries

Mechanisms encoded by diverse sequences are often difficult to compare directly, as their regulatory logic may be organized around distinct motifs or contextual features. Global libraries provide one way to address this challenge by anchoring sequences to a fixed pattern (or set of patterns) embedded in randomized sequence contexts, analogous to sufficiency experiments used to test motifs and their syntax.[12] This design establishes a common reference point, allowing SEAM to cluster mechanistic variation around the chosen pattern(s) and to systematically examine how their function depends on sequence context (Supplementary Fig. 22a).

We applied SEAM to global libraries containing CREB/ATF binding sites (`TNNTGAAAT`, with variable positions at `N`) using DeepSTARR.[13] This analysis revealed a range of mechanistic trajectories associated with changes both within the TFBS core and in surrounding nucleotides (Fig. 22e). In a separate analysis, SEAM uncovered program-specific patterns of motif utilization when the same global library was analyzed in the context of attribution maps representing either developmental or housekeeping regulatory programs. In several cases, sequence elements were subtly repurposed to adapt to different regulatory logic—for example, a DRE motif characteristic of housekeeping programs was reinterpreted as a GATA motif in the developmental context (Fig. 22f). We also extended this approach to cropped genomic sequences aligned on AP-1 instances, enabling direct comparison between synthetic global libraries and natural genomic contexts (Supplementary Fig. 22b). While the detailed patterns differed, SEAM identified comparable mechanistic spectra in both settings, suggesting that global libraries can serve as a scaffold for mapping genomic instances in attribution space (Supplementary Fig. 22c,d).

Overall, these exploratory analyses indicate that SEAM can be extended to global library designs, where anchoring to fixed sequence patterns provides a common basis for comparing context-dependent mechanisms. While the present results are illustrative, they highlight the potential of global libraries to complement local mutagenesis and design-derived approaches by offering a systematic framework to study how motif identity and surrounding context jointly shape cis-regulatory mechanisms.

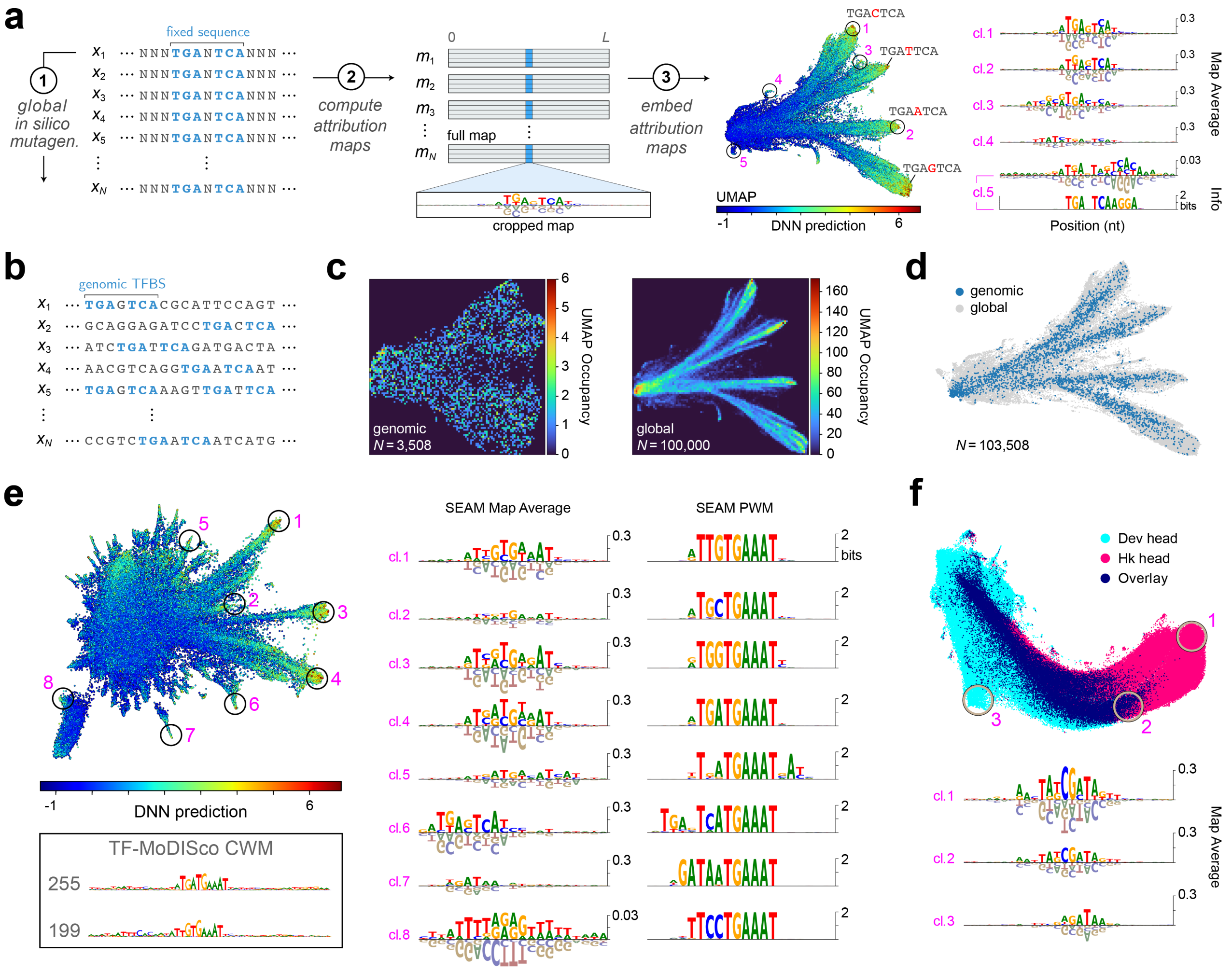


**Supplementary Figure 22. SEAM enables global analysis of motif-centric mechanisms across diverse sequence contexts a**, Schematic for applying the SEAM framework using global libraries to explore mechanistic variation of the AP-1 binding motif, using the fixed sequence `TGANTCA`. For every sequence in the global library, SEAM generates an attribution map that can be cropped to modulate the resolution of features obtained in the subsequent embedding. Attribution maps are then embedded to observe their spatial relationships. The maps within each encircled region in mechanism space are averaged to reveal noise-reduced attribution maps. **b**, Schematic for selecting all genomic instances of the AP-1 TFBS from the DeepSTARR test set. Genomic-based attribution maps for all 3,508 genomic instances were generated and cropped with 10 flanking nucleotides on each side of the AP-1 site, matching the length used in the global library. **c**, Left: Histogram of the UMAP embedding of the genomic-based attribution maps. Right: Histogram of UMAP embedding of global-based attribution maps, showing a substantially higher resolution of spatial features. **d**, UMAP embedding of both the genomic- and global-based attribution maps. **e**, Global analysis of CREB/ATF binding mechanisms using the sequence TNNTGAAAT (based on DeepSTARR's developmental head). Top left: UMAP embedding of cropped attribution maps. Bottom left: Sequence logo for previously published contribution weight matrices (CWM) derived by TF-MoDISco,[4] highlighting two distinct CREB/ATF motifs with 255 and 199 supporting seqlets, respectively. Right: Sequence logos for SEAM-derived averaged attribution maps for various clusters and associated sequence logo for the sequences within the cluster defined by the encircled regions in the UMAP embedding. **f**, UMAP embedding comparing attribution maps from the DRE global library (fixed sequence TATCGATA) generated using the DeepSTARR developmental (Dev) and housekeeping (Hk) heads.

### References

1. Gupta, H. *et al.* Highly diversified core promoters in the human genome and their effects on gene expression and disease predisposition. *BMC Genomics* **21**, 842, DOI: 10.1186/s12864-020-07222-5 (2020).
2. Rice, P., Longden, I. & Bleasby, A. EMBOSS: The european molecular biology open software suite. *Trends Genet.* **16**, 276–277, DOI: 10.1016/S0168-9525(00)02024-2 (2000).
3. Crutchfield, J. P. & van Nimwegen, E. The evolutionary unfolding of complexity. In Landweber, L. F. & Winfree, E. (eds.) *Evolution as Computation*, 67–94 (Springer Berlin Heidelberg, Berlin, Heidelberg, 2002).
4. Shrikumar, A. *et al.* Technical note on transcription factor motif discovery from importance scores (TF-MoDISco) version 0.5.6.5 (2020). 1811.00416.
5. Smilkov, D., Thorat, N., Kim, B., Viégas, F. & Wattenberg, M. SmoothGrad: Removing noise by adding noise. *arXiv* (2017).
6. Cochran, K. *et al.* Dissecting the cis-regulatory syntax of transcription initiation with deep learning. *bioRxiv* DOI: 10.1101/2024.05.28.596138 (2024).
7. Schreiber, J., Lu, Y. Y. & Noble, W. S. Ledidi: Designing genomic edits that induce functional activity. *bioRxiv* DOI: 10.1101/2020.05.21.109686 (2020). https://www.biorxiv.org/content/early/2020/05/25/2020.05.21.109686.full.pdf.
8. Zhou, J. & Troyanskaya, O. G. Predicting effects of noncoding variants with deep learning-based sequence model. *Nat. Methods* **12**, 931–934 (2015).
9. Sundararajan, M., Taly, A. & Yan, Q. Axiomatic attribution for deep networks. *arXiv* (2017).
10. Lundberg, S. M. & Lee, S.-I. A unified approach to interpreting model predictions. In *Advances in Neural Information Processing Systems*, vol. 30 (Curran Associates, Inc., 2017).
11. Bilodeau, B., Jaques, N., Koh, P. W. & Kim, B. Impossibility theorems for feature attribution. *Proc. Natl. Acad. Sci.* **121**, e2304406120, DOI: 10.1073/pnas.2304406120 (2024).
12. Koo, P. K., Majdandzic, A., Ploenzke, M., Anand, P. & Paul, S. B. Global importance analysis: An interpretability method to quantify importance of genomic features in deep neural networks. *PLoS Comput. Biol.* **17**, e1008925 (2021).
13. de Almeida, B. P., Reiter, F., Pagani, M. & Stark, A. DeepSTARR predicts enhancer activity from DNA sequence and enables the de novo design of synthetic enhancers. *Nat. Genet.* **54**, 613–624 (2022).
